# The extracellular milieu of *Toxoplasma*’s lytic cycle drives lab-adaptation and promotes changes in lipid metabolism primarily driven by transcriptional reprogramming

**DOI:** 10.1101/2021.03.09.434570

**Authors:** Vincent A. Primo, Yasaman Rezvani, Andrew Farrell, Amir Vajdi, Gabor T. Marth, Kourosh Zarringhalam, Marc-Jan Gubbels

**Affiliations:** Department of Biology, Boston College, Chestnut Hill, Massachusetts, USA; Department of Mathematics, University of Mass. Boston, Boston, Massachusetts, USA; Department of Human Genetics and USTAR Center for Genetic Discovery, Eccles Institute of Human Genetics, University of Utah School of Medicine, Salt Lake City, Utah, USA

**Keywords:** *Toxoplasma*, lab adaptation, serial passaging, experimental evolution, evolve and resequencing, regression analysis, genotype-phenotype correlation

## Abstract

To map host-independent *in vitro* virulence traits of *Toxoplasma gondii*, evolve and resequencing (E&R) during the lab-adaption was applied. Phenotypic assessments of the lytic cycle revealed that only traits needed in the extracellular milieu evolved. Surprisingly, only non-synonymous mutations in a P4 flippase fixed in two populations. However, dramatic changes in the transcriptional signature of extracellular parasites revealed a “pro-tachyzoite” profile as well as upregulation of fatty acid biosynthesis (FASII) pathway genes. More general, a set of 300 genes which expression profile changes during evolution mapped to specific traits. Validation of a select number of genes in this set by knock-outs indeed confirmed their role in lab-adaptation. Finally, assembly of an ApiAP2 and Myb transcription factor network revealed the transcriptional program underlying the adapting extracellular state. Overall, E&R is a new genomic tool successfully applied to map the development of polygenic traits underlying *in vitro* virulence of *T. gondii*.

## Introduction

*Toxoplasma gondii* is an apicomplexan parasite able to infect virtually any warm-blooded animal and causes opportunistic infections in humans [1]. Disease manifestations are typically mild, but severe and life-threatening illness is associated either with low immunocompetence, or the result of specific parasite and host genotype combinations leading to failures in immune response. For example, the right combination of ROP5, ROP17 and ROP18 alleles enables mouse immune response evasion, but not the human response [2–4]. Disease severity is also defined by additional host-independent virulence traits which define aspects of the lytic cycle such as replication rate, host cell invasion capacity, tissue transmigration efficiency, and enhanced survival in the extracellular environment [5]. Identifying the genetic basis of these host-independent virulence traits will provide insights in universal virulence mechanisms.

During *in vitro* lab-adaptation (i.e. from *in vivo* isolation to continuous *in vitro* cultivation) many of the general, host-independent virulence traits become enhanced [5, 6]. The lab-adapted RH strain used for most *in vitro* studies produces >5-fold bigger plaques *in vitro* (i.e. *in vitro* virulence) compared to several non-lab-adapted strains of the same Type I genotype [5]. In particular, RH’s replication rate and extracellular viability are superior to the GT1 strain, which is a non-lab-adapted Type I strain used for *T. gondii’s* reference genome [5, 7]. Efforts to identify the genetic basis for these phenotypic differences identified 1,394 single nucleotide polymorphisms (SNPs) between RH and GT1, of which 133 caused nonsynonymous amino acid changes and 54 were insertions/deletions (indels) within predicted coding regions [8]. Since experimental validation by allele swapping experiments did not reveal major drivers of RH’s *in vitro* virulence [8] we hypothesize that the genetic basis is either a combination of alleles (i.e. epistasis) [8]. However, the limited chronological record of RH’s *in vitro* history prevents dissection of genotype-phenotype development during the lab-adaptation process.

Evolve and resequencing (E&R) is a universal tool to dissect the genetic basis of adaptive or selective processes as it permits real-time investigation of genetic factors underlying experimental evolution [9, 10]. The most famous is Lenski’s Long-Term Experimental Evolution (LTEE) experiment of *Escherichia coli* [11] which identified alleles and expression profiles responsible for adaptation [12–16]. We applied E&R to the non-lab-adapted *T. gondii* GT1 strain to both establish a chronological record of *T. gondii* lab-adaptation and identify the genes and mechanisms underlying *T. gondii*’s host-independent, *in vitro* virulence traits. Over two years of lab-adaptation we observed augmentation of extracellular survival and host cell invasion capabilities, but no changes in intracellular replication rate. Despite a steady phenotype adaptation rate, only one gene with a non-synonymous SNPs fixating in evolving populations. Instead, the genetic basis of lab adaptation was found in ∼1000 differentially expressed genes (DEGs), almost exclusively restricted to extracellular parasites. The transcriptional signature revealed shut-downs of differentiation exits resulting in a “pro-tachyzoite” profile as well as upregulation of almost all genes of the fatty acid biosynthesis (FASII) pathway. Advanced DEG regression and clustering analysis identified 300 DEG’s strongly correlating with changes in phenotypes. Many of these genes are hypothetical but experimental validation of select genes confirmed their roles in GT1’s evolution. Finally, the transcriptional data were used to assemble a transcriptional network, which consolidated the defined roles of several characterized transcription factors [17, 18]. Taken together, we successfully applied lab-adaptation and E&R as a tool to identify a rich set of 300 host-independent virulence factors.

## Results

### Lab-adaptation of GT1 results in enhanced *in vitro* virulence

To ascertain a homogenous starting population, we first established several independent clones from a limitedly passaged, the cryopreserved GT1 strain used to establish *T. gondii*’s reference genome [7]. We propagated the original un-cloned GT1 strain (designated B0) and three independent GT1 clones (B2, B4, B6) in immortalized human foreskin fibroblasts (HFFs). Five percent of the parasite culture was passaged every 2-3 days for up to 223 serial passages (P) spanning ∼2 years and about 1500 parasite generations (Figure 1a).

**Figure 1.**
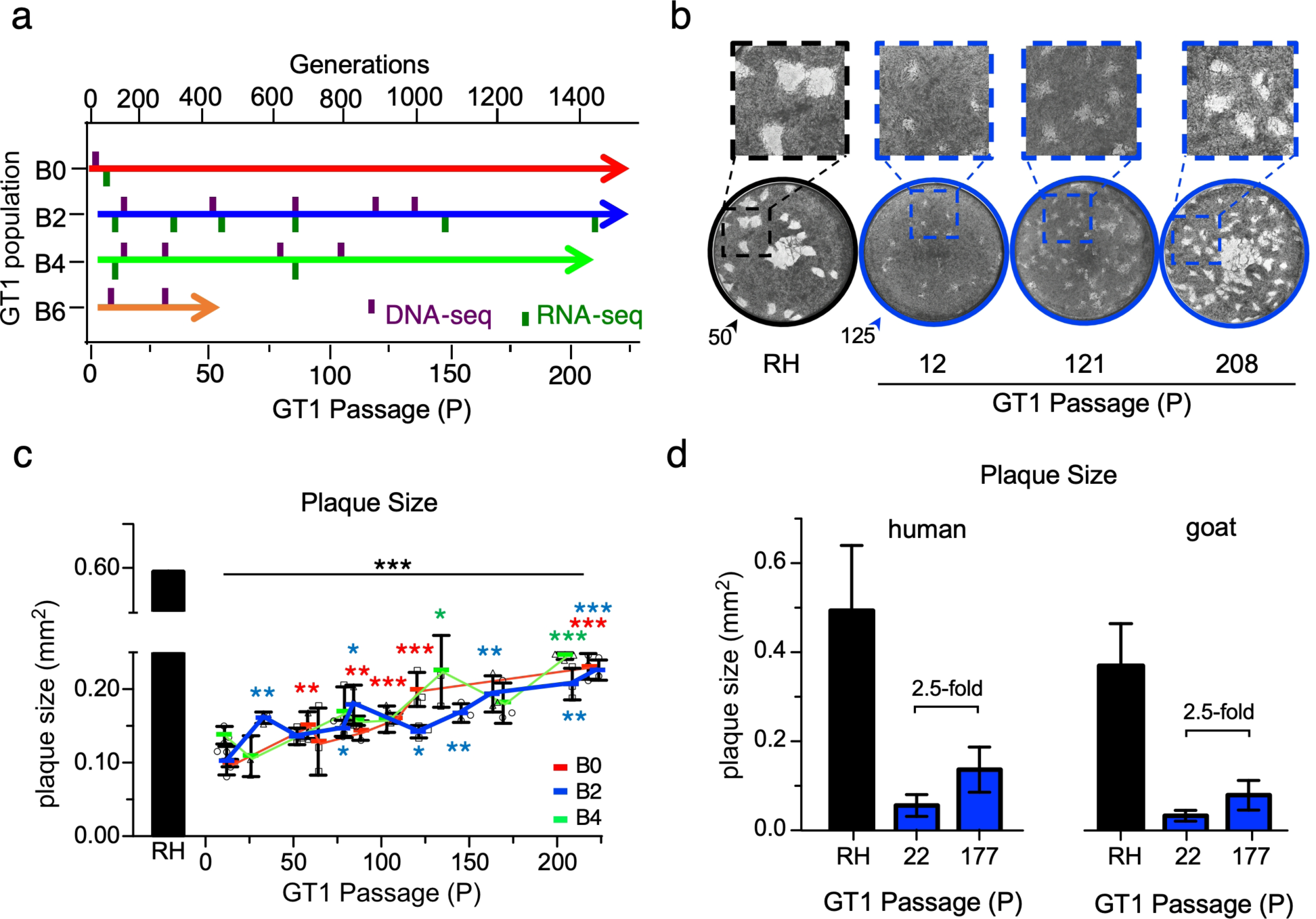
Augmented *in vitro* virulence of GT1 following lab-adaptation. **a** Experimental timeline of parallel GT1 lab-adaptation experiments and DNA-Seq/RNA-Seq timepoints, drawn to scale. B0 represents the polyclonal starting line, whereas B2, B4 and B6 represent distinct clones generated from B0 at the lowest possible passage. **b** Representative images of an 11-day plaque assay on HFF host cells. Arrow heads indicate the number of RH or GT1 (B2) parasites inoculated. **c** Quantification of plaque area following 11-day plaque assays with RH or GT1 (B0, B2, B4; color coded as in panel a) parasites. Black asterisks (*) indicate the *p-*value of the indicated GT1 passages relative to RH; colored asterisks corresponding with the lineages (*, *, *) indicate the *p-*value of the indicated GT1 passage relative to the respective population’s earliest passage. * = *p*-value *≤* 0.05, ** = *p*-value *≤* 0.01, *** = *p-*value *≤* 0.001. Colored blocks indicate mean of *≥*3 biological replicates for B0, B2, B4 (shown as squares, circles, and triangles, respectively) with error bars representing SD. One biological replicate is the mean quantification of *≥*25 plaques. **d** Plaque assay of RH or GT1 (B2) using both human-(HFF: human foreskin fibroblasts) and goat-derived (GSF: goat skeletal fibroblasts) host cells; n= 1, error bar indicates SD of *≥*20 plaques across two technical replicates.

As a proxy for GT1’s lab-adaptation we performed plaque assays along the evolutionary path (Figure 1b). Plaque size increased steadily: between first and last measurements, B0 increased 2.30-fold (*p*-value *≤*2.3*10^-4^), B2 increased 2.19-fold (*p*-value *≤*8.9*10^-4^) and B4 increased 1.86-fold (*p*-value *≤*1.1*10^-4^) (Figure 1c). However, plaque size of the lab-adapted RH strain, which is used as golden standard throughout our experiments, remained 2.31-to 2.62-fold larger than all >P200 GT1 populations (*p*-value *≤*6.13*10^-5^), indicating that after 2 years *in vitro,* GT1 has not yet reached the adaptation level of RH (Figure 1c).

Because RH and GT1 strains were isolated from different species, human and goat, respectively, we tested whether the difference in *in vitro* virulence was associated with the original host. We therefore compared plaque sizes of RH, B2-P22, and B2-P177 in primary HFFs with goat skeletal fibroblasts (GSF). This revealed that our lab-adaptation was host species independent (Figure 1d).

### Lab-adaptation of GT1 results in enhanced extracellular survival and invasion capacities

To define which step(s) in the lytic cycle contribute to the increase in plaque size we performed several functional assays. We first tested the extracellular survival capacity. GT1 passages were subjected to extracellular conditions for 0-10 hrs and viability of the population assessed hourly by plaque assay, resulting in survival curves (Figure 2a). The lethal time to kill 50% of the input population (LT50) was only 2 hrs for B2-P13, but increased to 5 hrs by B2-P211 (*p*-value ≤1.3*10^-4^) (Figure 2a). Measuring the area under these survival curves indicated that B2’s overall extracellular viability during the 10 hr exposure increased 2.21-fold by B2-P211, relative to B2-P13 (*p*-value *≤*2.5*10^-3^) (Figure 2b). Statistically, the LT50 of the B2 was already statistically similar to the well adapted RH reference strain by B2-P50, suggesting this is a very fast adapting feature.

**Figure 2.**
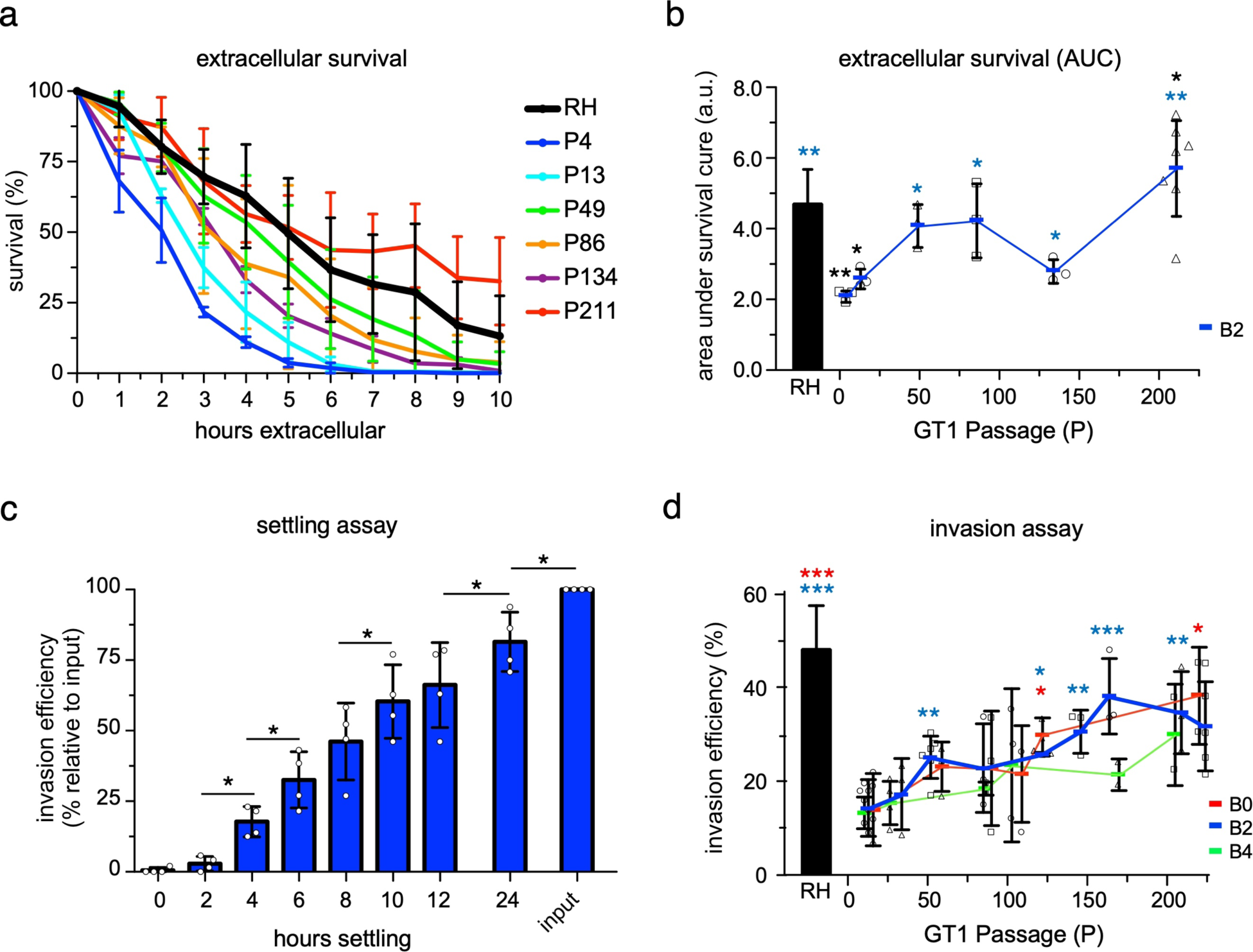
Lab-adaptation augments virulence traits of the extracellular milieu of the lytic cycle. **a** Mechanically released RH and GT1 parasites (B2) were incubated without host cells for 0-10 hrs and survivability was measured hourly by plaque assay. Colored block indicates mean of *≥*3 biological replicates with SD plotted; each biological replicate comprises 2 technical replicates. **b** Plot of the area under the survival curves shown in panel a. Blue block indicates mean of *≥*3 biological replicates (shown as squares, circles, and triangles) with error bars representing SD. **c** Mechanically egressed RH parasites were allowed to settle by gravity onto host cells for the times indicated before washing away extracellular parasites and analyzing invasion efficiency by plaque assay; data is normalized to an unwashed, total inoculum control (input). Mean of 4 biological replicates (circles) with SD is plotted; each biological replicate comprises 2 technical replicates. **d** Invasion efficiency: quantification of the total number of plaques formed relative to input following mechanical egress of RH or GT1 (B0, B2, B4) parasites. Horizontal lines indicate mean of biological replicates for B0 (n= *≥*3), B2 (n= *≥*3), B4 (n= 2), shown as squares, circles, and triangles, respectively, with error bars representing SD. For all panels: black asterisks (*) indicate the *p-*value of the indicated GT1 passage relative to RH; colored asterisks (*, *) indicates the *p-*value of the indicated GT1 passage relative to the respective population’s earliest passage; * = *p-*value *≤* 0.05, ** = *p-*value *≤* 0.01, *** = *p-*value *≤* 0.001;

We wondered how and why lab-adaptation could induce such a strong selection pressure onto the parasites. We reasoned that during the 1:20 serial passaging, the inoculated parasites might spend significant time extracellularly while settling by gravity onto the new host monolayer. To test this hypothesis, we determined the kinetics of this process by plaque assay using RH parasites. Indeed, only 50% of RH parasites successfully infected a new T25 host cell monolayer after 8 hrs and not even all parasites with plaque forming capacity had settled after 24 hrs (Figure 2c). Thus, extracellular survival is a strong selection pressure during our *in vitro* lab-adaptation protocol.

Next we tested host cell invasion efficiency, measured as the number of plaques relative to inoculum size. Time spent extracellularly was standardized across all samples by mechanically releasing the parasites from their host cells by needle passage. All GT1 populations showed a steady increase in invasion efficiency between the first and last measuring point: B0 increased 2.76-fold (*p*-value *≤*0.03), B2 2.24-fold (*p*-value *≤*0.003), and B4 2.27-fold (n=2; no statistics performed) (Figure 2d). Interestingly, invasion efficiency of RH remained 2.24-to 2.77-fold larger than all >P200 GT1 populations (*p*-value *≤*0.007), suggesting that continued lab-adaptation of GT1 will likely result in a continued rise of this virulence trait (Figure 2d). Hence, GT1’s invasion capacity is a lab-adaptive virulence trait.

We assessed the replication rate by enumerating the parasites per vacuole after 24 hrs of replication. We only observed only very minor (up and down) shifts in replication efficiency or doubling rates of B2 or B4 populations during lab adaptation (Supplemental Figure 1a-c). To complete the lytic cycle, egress efficiency was assessed on B2-P12, B2-P83, and B2-P211 by triggering with Ca^2+^ ionophore A23187 or ethanol (Supplemental Figure 1d). We observed no significant differences egress capacity. Thus, neither replication rate, nor egress efficiency are lab-adaptive traits in our study. In conclusion, traits facing the extracellular milieu of the lytic cycle are positively selected while traits within the intracellular milieu are not.

### WGS identified P4-flippase as a candidate polymorphic virulence factor

Short-read whole-genome sequencing (WGS) was performed with the GT1 populations at several passages to track genomic mutations during evolution (Figure 1a). We first assessed the clonality of three clonally-derived populations (B2-P15, B4-P15, and B6-P10) and compared it to the polyclonal the parent population (B0-P4). No mutational differences were identified between any of these low passage strains, indicating strong clonality within all of our starting GT1 populations.

In higher GT1 passages of B2, B4, and B6 we mapped many high-quality polymorphisms across all three evolving GT1 populations (Figure 3 & Supplemental Figure 2). B4 was sequenced at P15, P32, P79, and P105 spanning ∼800 generations, for which we observed two intronic mutations fixating in the population (Supplemental Figure 2a). Both are located within the 13^th^ intron of a dynein heavy chain (TGGT1_203135) and became fixed by P32. No non-synonymous mutation fixated in the B4 population. The evolving B6 population was sequenced only at P10 and P33, spanning ∼300 generations (Figure 1a). The B2 population was sequenced at P15, P52, P86, P120, and P135, spanning ∼1000 generations (Figure 1a). Only one non-synonymous mutation, L270R, emerged in a phospholipid-translocating P-type ATPase (P4-flippase) gene (TGGT1_245510) and remained fixed within the evolving population (Figure 3a). PCR plus Sanger sequencing revealed the L270R mutation as early as B2-P20 and fixating within the population by B2-P32 (Figure 3a). For B6, one non-synonymous mutation, A477D, within the same P4-flippase gene fixated within the population by P33 (Figure 3b). To gauge the genetic complexity of GT1 during lab-adaptation we sequenced five clones derived from B2-P86 (Supplemental Figure 2b). 17 mutations were uniquely mapped in single clones; the P4 flippase L270R mutation was shared across all clones. Although six of the mutations were non-synonymous, none fixated in the population, suggesting no fitness advantages were conferred. However, the complex population structure supports the random accumulation of mutations during evolution.

**Figure 3.**
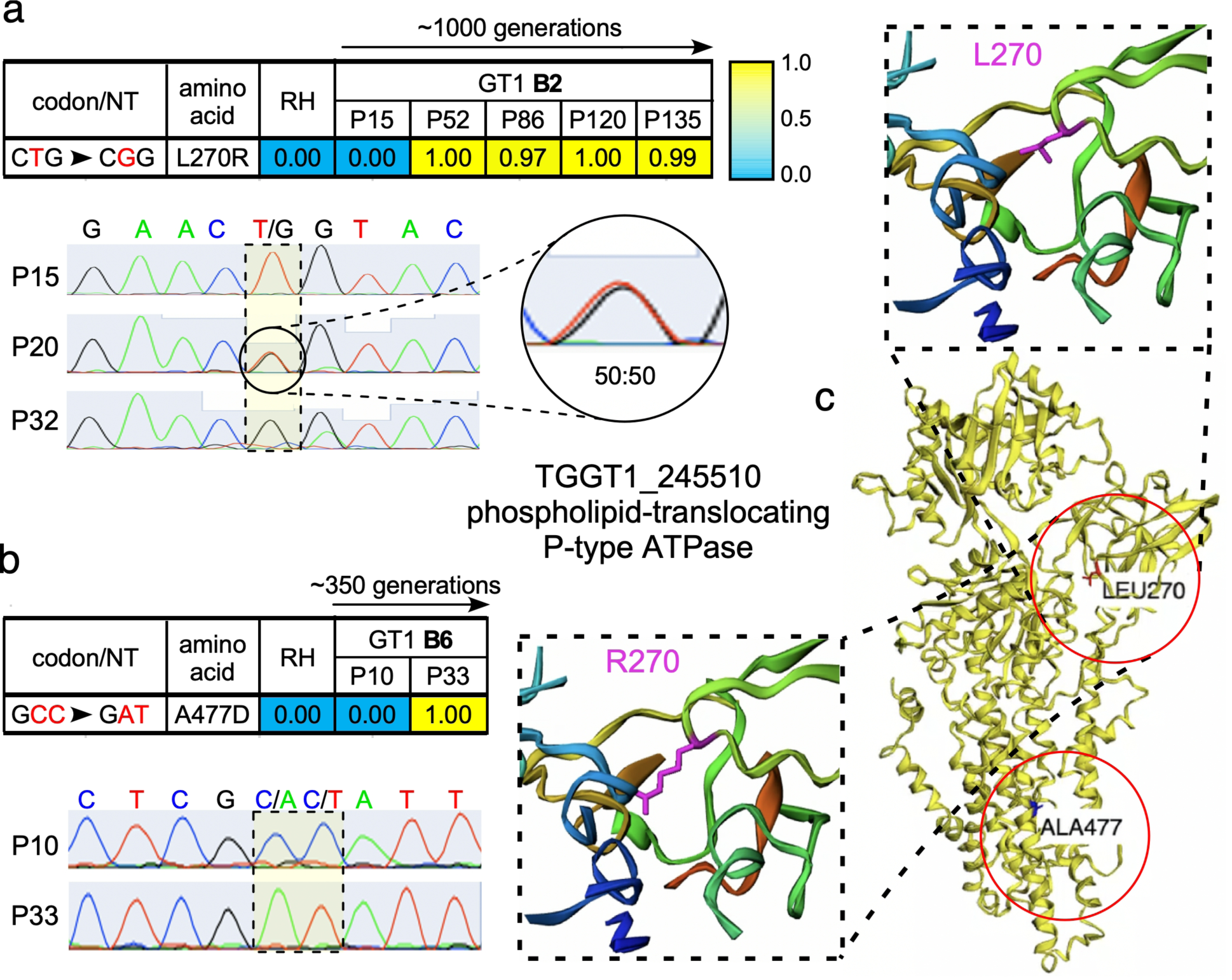
Lab-adaptation identified genomic mutations within a P4-flippase gene. WGSS identified the emergence of indicated mutations with the indicated allele frequency in GT1 populations; RH being shown for reference. **a** High-frequency (*≥*0.75) mutations identified in GT1 population B2 or **b** GT1 population B6. Top section: allele frequency represents the percentage of reads with the indicated allele. Lower section: Chromatogram of Sanger-sequenced PCR products confirms the presence of the P4 flippase L270R and A477D mutations in GT1 populations B2 and B6, respectively. **c** Structural predication of the P4 flippase protein and mapping of both mutations. Zoomed in images of the L270 and R270 alleles reveals the difference in sidechain extension into cavity space.

The identification of two different non-synonymous mutations in the same gene across two parallel evolving lines strongly suggests that these changes confer critical fitness benefits during lab-adaptation [16, 19]. To gauge potential effects on the function of the P4 flippase we modeled the *Toxoplasma* gene using Phyre2 [20] and predicted the impact of the SNPs on the structure using Missense3D predictive structural analysis [21]. The L270R mutation mapped to the cytoplasmic actuator (A) domain, which is responsible for inducing the functional E2-to-E1 conformational change by dephosphorylating the neighboring P domain [22]. The mutant allele results in a 78Å^3^ decrease in local cavity space due to the longer side chain of arginine (Figure 3c, enlargements). The A477D mutation is within the *α*-helix of the ATPase transmembrane (M) domain. It is therefore conceivable that these mutations affect either the efficiency or localization of the flippase protein, albeit further functional analysis is required.

### Lab-adaptation of GT1 results in few transcriptomic changes in the intracellular parasite

The single genomic mutation fixating in the population does not track with the continuing increase in plaque size during GT1’s lab-adaptation. Therefore, we reasoned that transcriptomic changes might be an alternative mechanism. To this end we performed mRNA-Seq (RNA-Seq) on asynchronously replicating intracellular GT1 B2 parasites at passages P11, P84, and P148. Read alignment and transcript estimation with HISAT2 [23] and featureCounts [24], respectively, was applied to all sequenced samples to get the read counts used for downstream analysis (Figure 4). Differential expression analysis (DEA), of P84 and P148 relative to the earliest timepoint, B2-P11, only identified 12 DEGs with many genes (9/12) annotated as hypothetical (Supplemental Table 1). The limited number of DEGs indicated that the intracellular state of GT1’s lytic cycle is likely not affected by lab-adaptation, which is corroborated by our phenotypic analysis of intracellular virulence traits.

**Figure 4.**
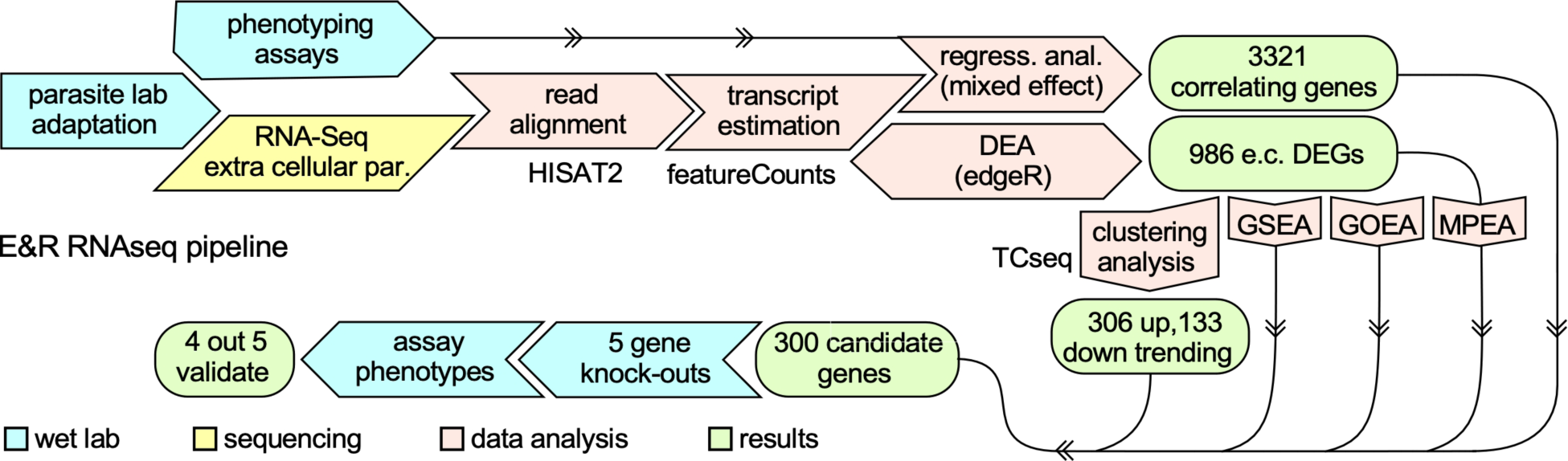
RNA-sequence analysis pipeline. Following lab-adaptation, 6-hour extracellular GT1 parasites at various timepoints were subject to short-read mRNA sequencing (Illumina). Read alignment with HISAT2 and transcript estimation with featureCount preceded regression analysis (RA) using a mixed-effect model and differential expression analysis (DEA) with edgeR. RA identified 3321 phenotype-correlating genes. DEA identified 988 differentially expressed genes (DEGs), relative to the earliest passage GT1 clone (B2-P11). Clustering the detected DEGs with time-course sequencing (TCseq) identified up- and down-trending gene clusters. The overlap of genes identified by DEA, TCseq, and RA identified 300 genes to be differentially expressed, correlating with at least one of the phenotypes, and trending up or down with the passage of time, thereby serving as candidate *in vitro* virulence factors that will likely provide additional insights into GT1’s extracellular adaptation. Gene Set-, Gene Ontology-, and Metabolic Pathway Enrichment Analysis (GSEA, GOEA, MPEA, respectively) along with life stage analysis, characterized the 986 DEGs and provided biological insights. Five representative upregulated genes (with a neutral fitness effect on the lytic cycle [31]) were selected for validation of their phenotype-conferring capacity by generating knock-outs in a high passage B2 strain.

### Lab-adaptation of GT1 is associated with many pro-tachyzoite, transcriptomic changes in extracellular parasites

Given the more prominent phenotype adaptations in extracellular parasites we hypothesized that differential gene expression might be associated with this state. RNA-Seq, transcript estimation and DEA on 6-hr extracellular B2 GT1 populations at passages P11, P35, P55, P85, P148, and P210 identified 986 significant DEGs (FC *≥*2, q-val *≤* 0.05), relative to the earliest passage, P11 (Figure 4). Of those, 435 DEGs were upregulated and 551 DEGs were downregulated (Supplemental Table 2).

Nearly 55% of these DEGs are of unknown function, challenging our ability to interpret the biology of the entire data set. Regardless of gene annotations, previously published RNA-Seq datasets allowed us to characterize these DEGs in the context of four *T. gondii’s* life-cycle stages (tachyzoite, bradyzoite, merozoite, and sporozoite), [25–27]. To characterize the developmental state based on an overall transcriptomic signature, we first developed a scoring method for a gene query based on how such genes are differentially expressed across all four developmental life stages (Supplemental Figure 3a). We validated this method by scoring previously published tachyzoite-, bradyzoite-, and sporozoite-associated gene sets [28]. Indeed, each gene set only scored significantly (*p*-value *≤* 0.05, bootstrap analysis) to their respective life stage, validating our scoring approach (Supplemental Figure 3b-d). Applied to the DEG dataset, significant upregulation of tachyzoite-associated genes, and significant downregulation of merozoite- and sporozoite-associated genes was revealed (Figure 5a). This indicates that lab-adaptation leads to a generally more tachyzoite-like expression profile in extracellular parasites. Interestingly, the bradyzoite-associated gene repertoire was almost equally up- and down-regulated, suggesting that extracellular parasites are in a quasi-bradyzoite-like state, likely due to the stress of the extracellular environment.

**Figure 5.**
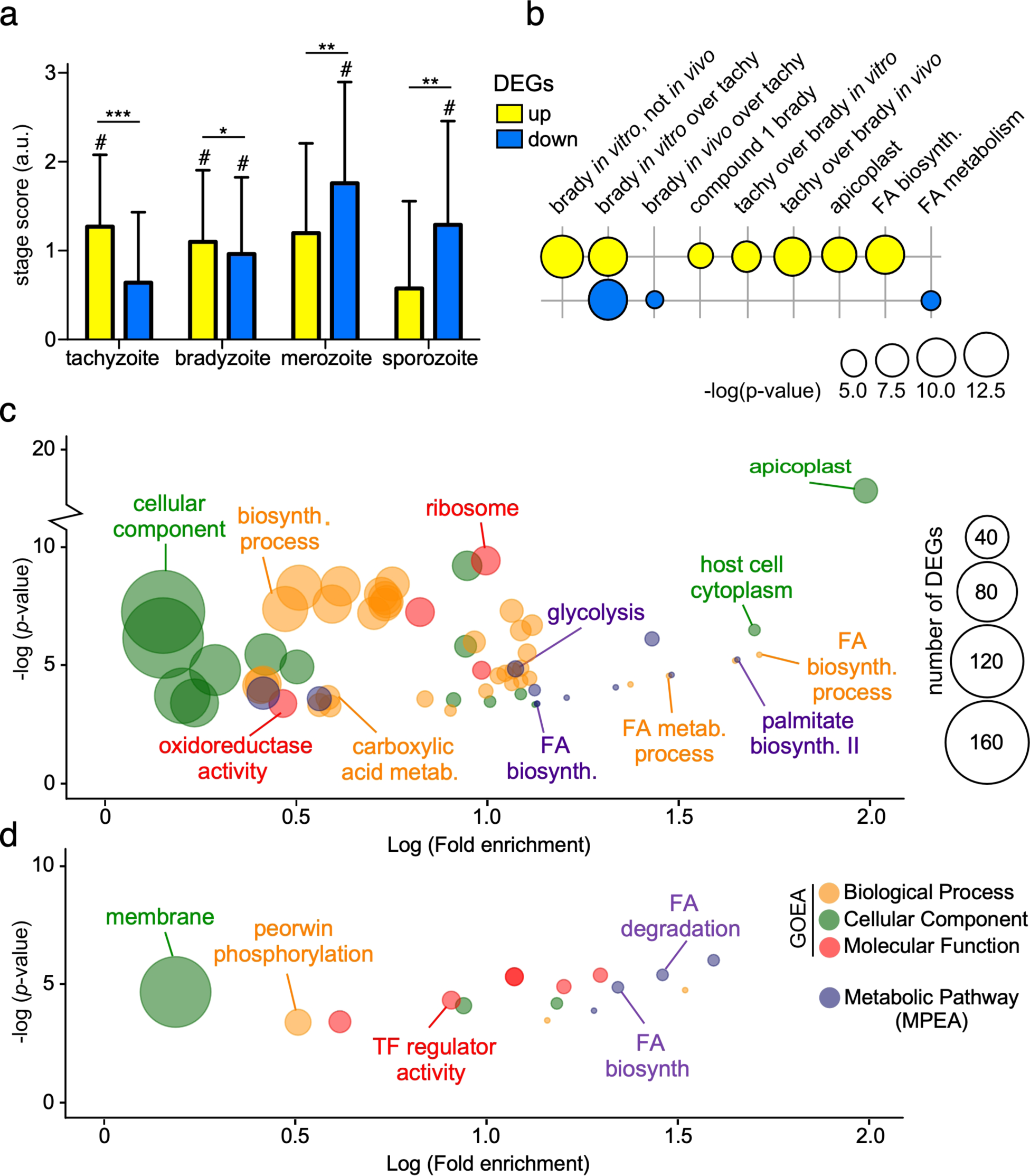
Biological insights of GT1’s evolved extracellular transcriptome. **a** Life stage analysis was utilized to characterize the types of life stage genes that are differentially expressed during GT1’s lab-adaptation; # indicate *p-*value *≤* 0.05, as determined by an independent bootstrap analysis (n=1000 random sampling) for each individual life stage; * = *p-*value <10^-8^, ** = *p-*value <10^-16^, *** = *p-*value <10^-32^ as determined by two-tailed *t*-test. **b** Gene set enrichment analysis (GSEA), as well as **c-d** GOEA and MPEA of up- and downregulated DEGs identified an enrichment in FA metabolism genes being largely upregulated in the apicoplast. Note: all significant GSEA, GOEA, and MPEA hits can be found in Supplemental Table 3.

### Gene enrichment analyses suggest fatty acid biosynthesis as a lab-adaptive biological process in the extracellular milieu of GT1’s lytic cycle

To distill biological insights from the 986 DEGs identified during lab-adaptation, we employed three types of statistical enrichment analyses outlined in Figure 4: 1) gene set enrichment analysis (GSEA [28]); 2) gene ontology enrichment analysis (GOEA); and 3) metabolic pathway enrichment analysis (MPEA) (Supplemental Table 3). The upregulation of genes in the apicoplast and the misregulation of fatty acid (FA) metabolism genes were in common across all three analyses: GSEA and GOEA identified the upregulation of FA biosynthesis while MPEA identified both up and downregulation of FA biosynthesis, as well as the downregulation of FA degradation (Figure 5b-d). Specifically, the majority of genes feeding into and within the apicoplast’s FASII pathway became upregulated during GT1’s lab-adaptation (Figure 6a-b). While short carbon chain FAs produced in the apicoplast can be further elongated in the endoplasmic reticulum (ER), we only observed a modest upregulation of the FA elongation pathway (Figure 6c-d). Taken together, these data suggest that FA availability is a selective pressure in the extracellular milieu.

**Figure 6.**
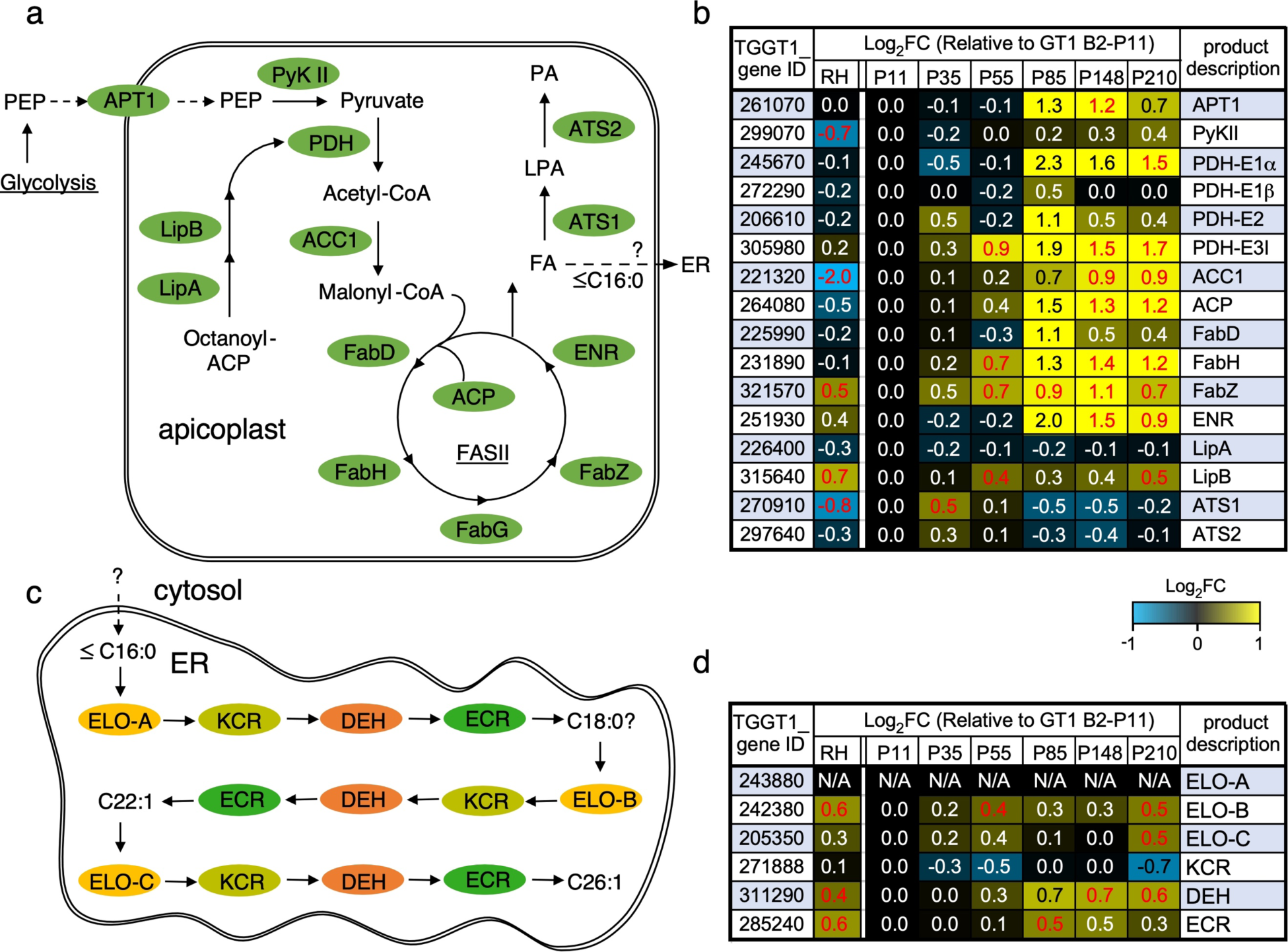
The FASII pathway becomes upregulated in extracellular GT1 during lab-adaptation. **a** Glycolysis produces PEP, which is transported into the apicoplast by APT1 and converted into pyruvate by PyKII [81]. Once lipoylated by LipA/B, the PDH complex converts pyruvate into acetyl-CoA, which is then metabolized to generate malonyl-CoA, the precursor metabolite required for the FASII pathway and FA synthesis [82, 83]. **b** Expression of the genes involved in the multi-step process of *de novo* fatty acid synthesis within the apicoplast. **c** Medium chain FA are translocated from the apicoplast to the ER for repeated rounds of carbon chain elongation by ELO A/B/C, KCR, DEH, and ECR [84]. **d** Expression of several genes involved in fatty acid elongation within the ER. Red text indicates *p*-value *≤* 0.05.

### Acquired *in vitro* virulence is a polygenic trait sustained by gene expression

The 986 identified suggest that lab-adaptive *in vitro* virulence is even a more multi-genic (and likely epigenetic) trait than we initially hypothesized. To further refine our list of 986 DEGs, we performed several analyses to calculate gene trends and correlation of genes with phenotypes as outlined in the following steps (Figure 4): 1) a linear mixed-effect regression model with smoothing B-splines was used to fit the time-course gene expression and phenotype data, 2) the inferred mean curves were sampled at regular intervals to align the phenotype and gene expression data and to calculate the correlation between each phenotype and gene expression curves, 3) gene expressions that strongly correlated (*R*^2^*≥*0.70, Spearman correlation) with GT1’s evolved phenotypes over time were identified, 4) a time-course clustering algorithm (TCseq; soft clustering with fuzzy cmeans) was employed to identify group of genes with similar pattern of expression over time and regression line was then fitted to gene clusters to quantify up and down-trending genes [29] (Supplemental Figure 4); 5) finally, the overlap of 986 DEGs that correlated highly with a phenotype (3321total) and trending genes (306 trend up; 138 trend down) were calculated resulting in a final list of 300 highest phenotype-correlating genes (Figure 4; Supplemental Table 4). We assembled the subcellular localizations for this gene set from the unbiased tachyzoite HyperLOPIT proteome analysis [30]. We could assign localization for 131 gene products, which consolidated the prominent role for the apicoplast, next to highlighting the secretory pathway as potential pressure for need of lipids (Figure 7a). Interestingly, the data covered 122 of the 193 upregulated genes but only 9 of the 107 downregulated genes, which makes this latter set rather enigmatic (also sparse by GOEA and MPEA Supplemental Figure 5; Supplemental Table 4). Finally, correlation of the 300 genes with the trends in phenotype evolution linked 204 genes with plaque size, which largely overlapped with the 275 genes correlating with invasion efficiency, whereas 31 correlated with extracellular survival (Figure 7b). To obtain experimental support for the phenotype-correlating genes as potential *in vitro* virulence factors we selected five genes spanning across various fold changes and phenotypic associations for genetic disruption (Figure 7b, colored stars). Exclusively up trending genes were selected, which grants us the ability to knock out (KO) genes in high passage GT1 with minimal effects of lab-adaptation during the time it take to isolate the mutant. In addition, we focused on genes with a neutral fitness conferring effect (i.e. fitness score) as identified in the genome-wide CRISPR screen [31] in order to avoid KO of essential genes (Figure 7c). We selected a glycosyltransferase (Gnt1; TGGT1_315885, a E3-ubiquitin ligase [32]), motor protein myosin I (MyoI: TGGT1_230980, which resides in the residual body (RB) of intracellular parasites and facilitates cell-cell communication during division [33]), microneme protein 13 (MIC13; TGGT1_260190,which has been associated with oxidative stress survival through and unknown mechanism [34]) and two hypothetical genes (TGGT1_262590, TGGT1_264240) (Figure 7c).

**Figure 7.**
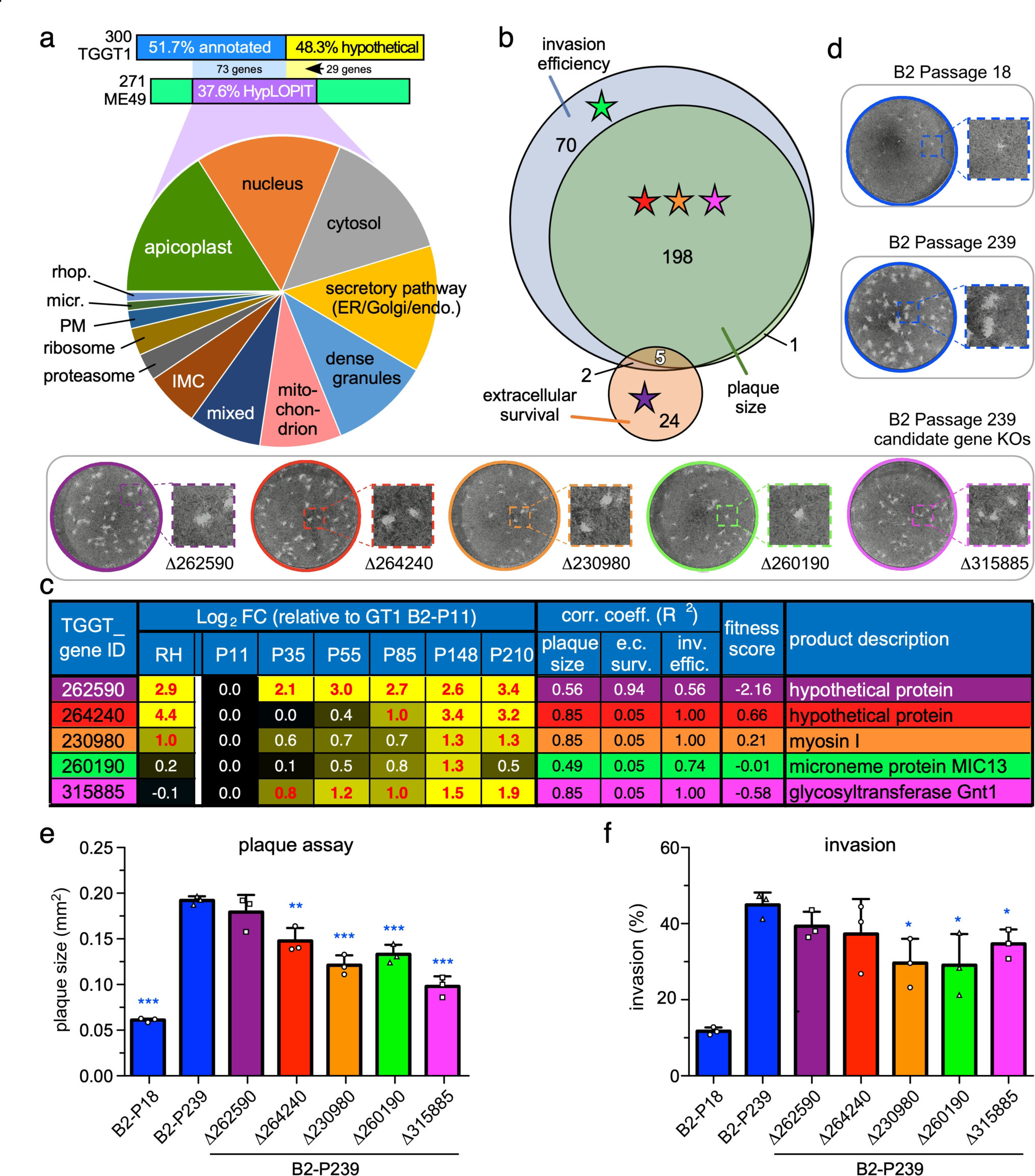
Functional analysis of candidate gene knock-outs identified several differentially expressed genes important for optimal *in vitro* virulence, suggesting acquired *in vitro* virulence is a polygenic trait. **a** HyperLOPIT sub-cellular localization data [30] on the 300 strongest phenotype conferring genes. Only 271/300 genes are cross annotated between the TGGT1 and TGME49 genome annotation (data is only available for the ME49 genes). 97 of the 107 genes with HyperLOPIT data are upregulated. The ‘mixed’ slice comprises genes with conflicting assignments between the two statistical algorithms used for the sub-cellular assignments. Data available in Supplementary Table 5. GOEA and GMEA analyses for this gene set is provided in Supplementary Figure 5. **b** Venn diagram of 300 strongest phenotype conferring genes and their correlation with lab-adaptive phenotypes. **c** Expression profiles (Log_2_-FC), phenotype correlation coefficients (R^2^), and fitness score in the genome wide CRISPR screen of the lytic cycle [31] of the five genes chosen for KO. Red text indicates *p*-value *≤* 0.05. **d-f** Upon successful KO (Supplementary Figure 6), plaquing capacity after 11-days (**e**) and invasion capacity (**f**) were evaluated. Mean of *≥*3 biological replicates (represented as squares, triangles, or circles) with SD is plotted. Blue asterisks (*) indicate the *p-*value of the indicated GT1 passages relative to B2-P239; * = *p*-value *≤* 0.05, ** = *p*-value *≤* 0.01, *** = *p-*value *≤* 0.001. For all panels, the five genetic KO’s are color coded.

The most evolved B2-P239 was used as parent for the gene KO experiments. Genotypes of isolated clones were confirmed by diagnostic PCR and the ablation of mRNA expression by qRT-PCR (Supplemental Figure 6). The five KO clones were evaluated for plaque size and invasion efficiency, using low passage B2-P18 and the B2-P239 parent as controls. Relative to B2-P239, four KO lines showed a reduction in plaque size (0.51-0.76-fold; *p*-value <0.01) (Figure 7d) while three lines displayed a reduced invasion efficiency (0.65-0.77-fold; *p*-value <0.05) (Figure 7e). Importantly, the KO parasites displayed the phenotypic characteristics predicted by their correlation coefficient (*R*^2^) (Figure 7b), indicating a high degree of accuracy in phenotype prediction from our Mixed-effect Regression Splines. Gene expression of MyoI and Gnt1 strongly correlated with plaque size and invasion efficiency (Figure7b-c), and KO of these genes validated their influence on those phenotypes (Figure 7d-f). On the other hand, TGGT1_262590 did not show strong correlation with plaque size and invasion efficiency during lab-adaptation (Figure 7b-c), and KO evaluation validated that relationship (Figure 7d-f). Based off each gene’s plaque size and invasion correlation coefficient (R^2^; Figure 7c), only two unexpected outcomes were observed: TGGT1_264240 displayed reduced invasion efficiency of ∼17% as expected by RA (*R*^2^ = 1.0), but this did not reach statistical significance (*p-*value = 0.24) due to the sizable standard deviation in our biological replicates (Figure 7f); the MIC13-KO displayed a significant ∼35% reduction in plaque size, which was much more dramatic than expected given its R^2^ value of only 0.49 (Figure 7c-d). Overall, the predicted phenotype for four out five genes was successfully confirmed. This strongly suggests that the list of 300 phenotype-correlating, trending genes (Supplemental Table 4) truly harbors many *in vitro* virulence factors, which indicates that GT1’s acquired *in vitro* virulence is a polygenic trait sustained by specific gene expression patterns.

### The transcriptional network of lab adaptation and the extracellular state

We sought to define the transcriptional network and transcription factors that orchestrate the transcriptomic changes of lab adaptation. The apicomplexan Apetala 2 (ApiAP2) family of transcription factors (TFs) is with 68 annotated members on ToxoDB the most expanded. Numerous biological traits have been associated with specific AP2s [17, 18, 35–40]. In addition, Myb TFs are another significant family (14 annotated genes on ToxoDB) associated with parasites specific functions such as BFD1 with bradyzoite differentiation [41]. We identified seven TFs whose expression in extracellular parasites significantly changed during lab adaptation by comparing expression level at B2-P11 with B2-P210 (Figure 8a; Supplemental Figure 7; note that AP2IX-9 is significant at P85 and P148 but not at P210). All six AP2 factors are trending down and the single up trending is a Myb TF, which we named Myb2 (TGGT1_306320). Two other Mybs (TGGT1_321450, TGGT1_264120) are highly expressed in extracellular parasites, although their pattern does not change (Supplemental Tables 2, 5). Interestingly, three down-trending AP2s, Ib-1, IV-3 and IX-9, are upregulated at the start of bradyzoite differentiation, but are downregulated again in mature bradyzoites [35]. Hence, this indicates that the overlap of DEGs between extracellular parasites and bradyzoites is also shared in their transcriptional regulation network.

**Figure 8.**
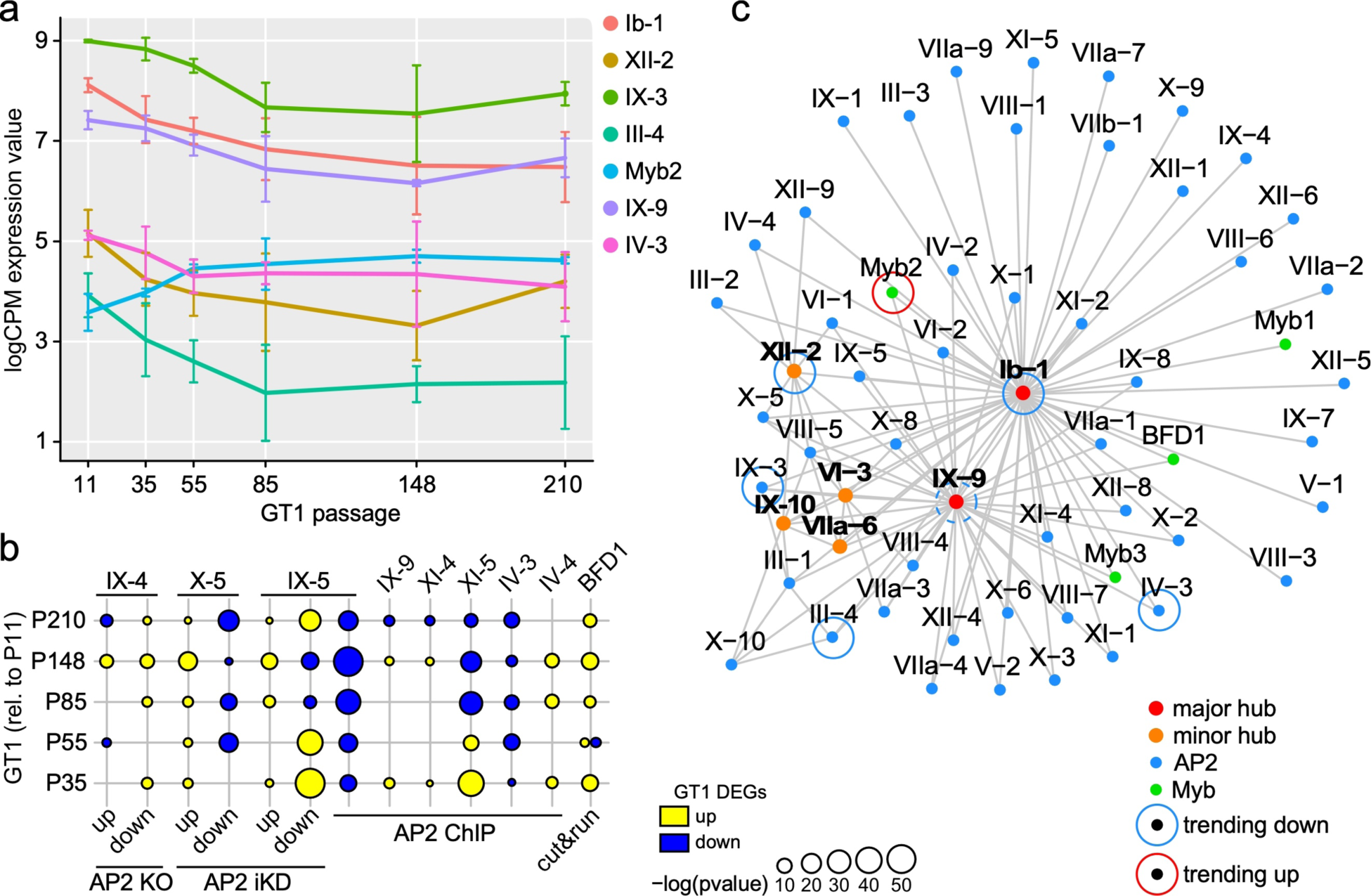
Gene expression network analysis with AP2 and Myb family transcription factors. **a** Inspecting the expression of AP2 and Myb TFs in extracellular parasites identified six down trending AP2s and one up trending Myb among which AP2Ib-1, AP2IX-3, AP2IV-3 and Myb2 representing statistically significant trend over the time based on R^2^ (*≥*0.5) of the regressed line to each of the TFs expression (Supplementary Figure 7) Though AP2IX-9 does not reach the statistical significance level of trending, it’s expression level is significantly changing between P11 and P148. Trends of the AP2s indicate their key role in extracellular transcriptional mechanism. **b** Previous studies have identified direct (by AP2 ChIP-Seq [17], ChIP-qPCR [35, 37, 39], ChIP-chip [40] or cut and run [41]) and indirect (by AP2 ikD or KO followed by RNA-seq [17, 18] or microarray analysis [38]) targets of AP2 and Myb TFs. GSEA comparing these direct/indirect transcription factor targets to lab-adaptive DEGs identified significant enrichments, indicating AP2 TFs as potential regulators of lab-adaptation. **c** Transcription factor expression network analysis incorporating both intracellular and extracellular RNA-Seq data over the evolutionary trajectory base on predicted AP2 and Myb transcription factors. The following Myb domain genes were considered; TGGT1_200385 (BDF1), TGGT1_203950, TGGT1_213890, TGGT1_264120 (Myb1), TGGT1_275480, TGGT1_306320 (Myb2), TGGT1_321450 (Myb3); TGGT1_215895; AP2IX-10 represents TGGT1_215895, which was not previously named.

The targets of APIV-3 and AP2IX-9 have been mapped [35] and we therefore intersected these with the DEGs during lab adaptation (Figure 8b). Indeed AP2IV-3 controlled genes trend down, but the effect is much weaker for AP2IX-9. AP2IX-9 expression peaks early in tissue cyst formation and actually suppresses it, while AP2IV-3 promotes this process and peaks later [35]. Lab adaptation therefore seems to be driven beyond the inhibition of ‘state change’ controlled by APIX-9. To be comprehensive, we included data for all TFs with known targets [17, 18, 35, 37-41]. Genes controlled by BFD1 are upregulated during lab adaptation, again consistent with genes shared with bradyzoite differentiation. Expression of most genes controlled by AP2XI-5 [40], AP2X-5 [18], and AP2IX-5 [17, 42] trends down. This is not surprising since these clusters associated with cell division, and extracellular parasites do not divide.

Finally, to comprehensively grasp any major epigenetic players of GT1’s polygenic lab-adaptation, we constructed a Protein-Protein Interaction (PPI) network of AP2 and Myb interactions using a Gaussian Graphical Model fitted on all RNA-Seq data points across passages from both intracellular and extracellular conditions (Figure 8c). AP2IX-9 and AP2Ib-1 appear as the main hubs in the transcriptional network: the former is a repressor and the latter a suspected activator of bradyzoite gene expression [35, 39, 43]. Moreover, the additional connection of BFD1 to these two main hubs fits with the shared bradyzoite profile. Furthermore, some of the interactions, such as the link between AP2IX-5 to AP2IX-9 and AP2XII-2, as well as the link between AP2X-5 and AP2Ib-1, have been previously reported and serve as calibration points of our analysis [17, 18]. Overall, our transcriptional network analysis confirms the significant overlaps with the bradyzoite differentiation process, but not with the mature bradyzoites, which fits with the mixed bradyzoite profile in the life stage analysis (Figure 5a). Moreover, AP2IX-9 and AP2Ib-1 are the main TFs associated with the extracellular state and both are key drivers of lab adaptation.

## Discussion

After ∼1500 generations of lab adaptation GT1 displayed significant evolution of its *in vitro,* host-independent virulence (reproducible >2-plaque size fold increase). However, plaque size is still >2-fold smaller compared to the lab-adapted RH strain. Indeed, the steady slope of plaque size increase (Figure 1c) indicates that at P200 GT1 is still on a continuing evolutionary path. Extracellular survival and invasion efficiency were the features most strongly evolved during lab adaptation, whereas changes in the intracellular replication cycle were not observed. Phenotype changes were driven by transcriptional reprogramming of extracellular parasites and we identified two biological drivers: 1. a shift toward a tachyzoite-enriched gene expression profile by shutting down genes expressed in other life stages that more closely resembles RH (Figure 5a; Supplemental Figures 5a); 2. enhanced *de novo* synthesis of FAs through upregulation of the FASII pathway (Figure 6). Germane to the former, the mechanism underlying increased *in vitro* growth rates in *Plasmodium* spp. is through shutting off sexual stages differentiation (through inactivation of gametocyte driving AP-G) [44, 45]. Hence, our finding fits this principle of preventing siphoning of lytic cycle parasites into different developmental stages. Although merozoite and sporozoite gene expression depleted during lab adaptation, bradyzoite genes were a mix of going up and down (Figure 5a-b). Analysis of the transcriptional network revealed that several TFs associated with early steps of bradyzoite differentiation [35] were upregulated during lab adaptation, which explains the upregulation of bradyzoite associated genes. We hypothesize that the downregulated portion is primarily associated with mature bradyzoites. The shared transcriptional network likely originates in stress, which is a key factor driving bradyzoite differentiation [46], and extracellular parasites are likely also under environmental stress as it is the leading evolving trait (Figure 2b).

Pertaining to the second biological pathway, the availability of FA in the extracellular milieu appears to be a limiting factor. Further support for a critical role for lipid homeostasis is provided by the SNPs in the flippase gene fixating early during lab-adaptation (Figure 3). P4-phospholipid flippases are transmembrane transporters of cations, heavy metals, and particularly, phospholipids across lipid bilayers [47]. Phospholipid flipping activity is of great biological importance for the biogenesis of vesicles [48, 49], creating a fusion-competent bilayer [50], maintaining membrane stability [51, 52], and generating signaling cues [53–55]. Indeed, secretory pathway genes also trend up during evolution (Figure 7a). It was recently reported this particular *Toxoplasma* P4-flippase localizes to the ER [56], while a whole parasite organelle proteomic approach assigned it to the a dynamic Golgi/PM localization [30]. Dissecting how the flippase SNPs modulate the kinetics or substrate preference will require extensive further work. A string of recent insights supports the role of the FASII pathway in *in vitro* virulence [57–60]. Furthermore, the availability of lipids in the extracellular environment has been identified as a critical factor for *in vitro* virulence in a dose-dependent manner [57, 61]. The highly lab adapted RH strain is able to maintain superior *in vitro* virulence even under lipid-starved conditions by upregulating its *de novo* FA/lipid synthesis up by 15% [57], which is supporting our data (although the flippase gene in RH contains no SNPs, suggesting an alternative path to the same state). Our lab adaptation experiment was performed under 1% FBS, which is a relatively low concentration and might make FAs a limiting factor. The critical insight regarding the contribution of lipids to *in vitro* virulence is the novel connection with surviving in the extracellular environment as the critical place of lipid availability.

The biology of *T. gondii* in the extracellular milieu has been largely understudied as it is regarded as a relatively short period bridging two intracellular replication cycles of low significance. However, ∼20% of tachyzoites in infected mouse tissue are extracellular, suggesting that tissue-residing parasites can spend significant time outside a host cell during their lytic cycle [62]. We demonstrated that parasite growth *in vitro* is inherently associated with a prolonged extracellular state. Thus, the extracellular tachyzoite is important both *in vivo* and *in vitro*. Transcriptomic studies report ∼2400 DEGs between parasites residing intracellularly vs. those in the extracellular environment, a huge portion of the ∼8000 annotated genes (Supplemental Figure 8a; Supplemental Table 6) [27, 63–65]. Since extracellular parasites are arrested in G0 of the cell cycle, we ascertained these DEGs are not merely the result of the stalled division cycle. Hereto we intersected this gene set with genes undergoing cyclical expression throughout the intracellular replication cycle (1967 total genes); This showed that ∼73% (1758 genes) of the extracellular DEGS do not overlap with the cyclic transcription patterns seen in the replication cycle (Supplemental Figure 8b). Taken together, the extracellular state is a transcriptionally and biologically unique but has been underappreciated for its selective pressure in lab adaptation.

Nearly half of the 300 genes with the highest lab adaptation correlation are hypothetical and lack protein localization data (90% of the downregulated genes fall in that class). This begs the question what do they do? Insights from the experimentally validated genes hints at two processes: up-regulation of MIC13 might be part of a stress response to the extracellular environment as indicated by a recent MIC13 study in RH growth under stressed conditions [34]. Second, studies on Gnt1 have shown that this glycosyltransferase incorporates GlcNAc onto Skp1 thereby promoting the formation of the E3-ubiquitin ligase containing SCF (Skp1/Cullin-1/F-box protein) complex [32, 66]. The SCF complex directs proteins for degradation by the 26S proteasome and is O_2_ regulated in *Dictyostelium* [67], indicating its role in maintaining redox homeostasis in the cell. Interestingly, during GT1’s lab-adaptation, six genes of the proteasome core complex were upregulated over time (Figure 7a; Supplemental Table 2), suggesting that lab-adaptation, results in increased protein turnover within extracellular parasites. Lastly, the identification of MyoI is quite peculiar as it resides in the residual body of intracellular parasites and is critical in maintaining parasite-to-parasite communication between dividing parasites [33]. However, extracellular parasites have no residual body, which suggests a novel function for MyoI. We will follow up on this new function in future work. Either way, further mining the genes associated with lab adaptation will map the nature of host independent virulence factors and likely reveal additional new biological insights.

To assure ourselves that absence of genomic mutations fixating in the population was not due to a low genomic mutation rate, we evaluated the genomic mutation rate throughout our experiment. The clones sequenced 71 passages apart (B2-P15 and five clones at B2-P86, Supplemental Figure 1b) were used to determine the mutation rate of GT1 at 1.1 x10^-10^ mutations/bp/generation (Supplemental Figure 9). This is within the range of RH’s reported mutation rate: 5.8 x10^-11^ mutations/bp/generation [68]. This rate predicts that 2-4% of a *T. gondii* population accumulates a single mutation within a single passage (2-3 days) under standard laboratory conditions. Thus, the lack of mutations fixating in the population is not due to an underpowered experimental design. Therefore, the lack of genomic mutation fixation is likely caused by the manifold of pressures on the switch to *in vitro* conditions. This might not permit fast selection for single traits, and instead is better addressed with more global changes in transcriptional programs. The pliable transcriptional network suggests that lab adaptation in our experiment is maintained epigenetically, which we will address in future work.

In conclusion, our results indicate that lab-adaptation of GT1 results in augmented phenotypes driven by selection pressures in the extracellular environment. Furthermore, our work demonstrates the phenotypic and transcriptomic versatility under laboratory conditions commonly practiced in the *T. gondii* field, which forewarns researchers to carefully consider their *in vitro* culture conditions (e.g. % FBS as FA source, settling time). The results also demonstrate the complex and polygenic nature of lab-adaptation and *in vitro* virulence. We have only scratched the surface of our 300 potentially phenotype conferring genes and therefore anticipate the discovery of many additional host-independent virulence factor in future validation work.

## Materials and Methods

### In vitro culturing of T. gondii

The GT1 strain of *T. gondii* was obtained through BEI Resources (catalog NR20728) and propagated into culture using ED1 medium [6] supplemented with 10 mM HEPES, pH 7.2. Parasites were maintained in human telomerase reverse transcription (hTERT) immortalized human foreskin fibroblasts in a humidified 37°C incubator under 5% CO_2_. Typically, early passage (P) GT1 parasites require 3-4 days to fully lyse a T25 (25 cm^2^) flask of host cells while later passage GT1 parasites (>P80) require 2-3 days. Passing was performed serially by transferring 500 µl of the lysed host cell flask (consisting of suspended parasites) into a new T25 flask of hTERT host cells containing 9 ml of warm ED1 media. Long 1 ml serological pipettes were used for transferring in order to reduce cross-contamination of separate *T. gondii* populations. Serial passaging of GT1 occurred in this fashion for up to 220 passages.

After successful establishment of the GT1 stabilate into culture, several single-cell clones derived from the initial population were obtained. Clonal GT1 populations, named “B1”, “B2”, “B3”, etc., as well as the original polyclonal population, named “B0”, were frequently frozen down during the ∼220 passages (∼2 years) of *in vitro* culturing. Frequent freeze downs of parasite populations ensured a chronologically maintained fossil record of the evolving parasites, as stored resource for future experiments.

### In vitro culturing of host cells

hTERT-immortalized HFF cells were maintained in T175 (175 cm^2^) flasks up until P18, upon which they are passed into T25 flask by P19 and used as hosts for parasite culture. Primary HFF cells are maintained in T175 (175 cm^2^) flasks up until P9, upon which they are passaged into desired flasks or plates by P10 and used as hosts for plaque assay or IFA. Goat skeletal fibroblast (GSF) cells were generously provided by Dr. Mahipal Singh of Fort Valley State University [69].

### Plaque assays

T25 flasks containing medium-to-large vacuoles of parasites (∼2 days post inoculation) were washed 2x10 ml with PBS to remove extracellular parasites and debris, followed by addition of 3-6 ml (volume dependent on vacuole size and number) of warm ED1 media. Next, the cell monolayer was scraped with a rubber police man and the host cells mechanically lysed by passing through a 27G needle. Mechanically egressed parasites were filtered through a 0.22 µm pore polystyrene filter to remove host cell debris. Parasite concentration was determined with a hemocytometer and adjusted to 10,000 cells/ml in a final volume of 3 ml Ed1 medium.

Three-to-six week-old primary HFF or GSF monolayers were used for plaque assays. Plaques for GT1 and RH were allowed to form for 11 days on primary HFFs or 14 days of GSF host cells. Host cell monolayers containing plaques were fixed with 100% ethanol for 10 min at RT, stained with 5X crystal violet solution for 10 minutes at RT, washed twice with PBS, and allowed to air dry for 24 hrs. Quantification of plaque size (i.e. *in vitro* virulence) and plaque number relative to input (i.e. invasion efficiency) was performed with FIJI [70].

### Extracellular survival assay

Parasite cultures were prepared as described above and 3 ml of the parasite cell suspension was incubated at 37℃ and 5% CO_2_ in non-tissue-culture-treated 6-well plates for 0-10 hours. Plaque assays were performed hourly and quantified as mentioned above. Plaque numbers at each timepoint were normalized to the 0-hour timepoint to yield percent survival.

### Replication assay

Mechanically egressed parasites (27G needle) were inoculated onto confluent primary HFF monolayers grown on coverslips in 6 well plates, centrifuged at 1000*g for 5 mins, allowed to invade at 37℃ (floating in a water bath) for 10 mins, and subsequently washed 3x with PBS. Intracellular parasites were then allowed to replicate for exactly 24 hrs followed by 100% methanol fixation and immunofluorescence assay (IFA) with rabbit *α*-TgIMC3 [71] to mark the cortical cytoskeleton and 4′,6-diamidino-2-phenylindole (DAPI) to mark DNA. The number of parasites per vacuole was enumerated for 100 vacuoles.

### Egress assay

Mechanically egressed parasites (27G needle) were inoculated onto 6-well HFF plates containing glass coverslips and allowed to invade and replicate for 30 hrs. Replacement medium containing either 1 μM of A23187, 5% ethanol, or DMSO was incubated for exactly five minutes in the plates before fixation of infected monolayers with 4% PFA and IFA with *α*-TgIMC3 and DAPI. The number of egressed vacuoles was enumerated for a total of 50 vacuoles per condition.

### DNA-sequencing and analysis

Parasite genomic DNA was isolated using Qiagen DNAeasy Blood and Tissue kit (catalog 69504) according to manufacturer’s protocol. Illumina’s Library Prep kit (FC-121-1030) was used to generate ∼361 bp DNA fragments, on average, which were quantified using Quibit Flex Fluorometer (catalog Q32851) and quality checked using Agilent’s TapeStation (catalog 5067-5584, 5067-5585). Next, 150 bp paired-end sequencing was performed on Illumina’s NextSeq500 platform using their high output flow cell kit (FC-404-2004) according to manufacturer’s protocol. FASTQ reads were then analyzed by RUFUS analysis to call sequence variants between two samples (https://github.com/jandrewrfarrell/RUFUS) [72, 73]. To compare RUFUS variants across all samples, all calls were merged and GRAPHITE was used to genotype calls across all samples in the study (https://github.com/dillonl/graphite). High frequency variants were called if the emerging mutation reached 75% allele frequency in at least one evolving population.

### P4-flippase genotyping

Parasite genomic DNA was isolated using Qiagen DNAeasy Blood and Tissue kit (catalog 69504) according to manufacturer’s protocol. M13 primers (Supplemental Table 7) used to PCR amplify ∼350bp region surrounding the R270L and A477D alleles. PCR products were purified and sent to Eton Biosciences for Sanger sequencing. Allele confirmation and chromatographs were obtained using 4Peaks (https://nucleobytes.com/4peaks/).

### RNA-sequencing and data analysis

#### Library Preparation and sequencing

*T. gondii* infected (24-36 hours) hTERT-immortalized HFF monolayers were washed 3x with PBS and either mechanically lysed with a 27 G needle (6-hour extracellular) or immediately lysed (intracellular) and processed on ice for RNA isolation using Qiagen RNAeasy kit (catalog 74104) according to manufacturer’s protocol. RNA quality was evaluated by measuring the RNA integrity Number (RIN) using Agilent’s TapeStation (kit catalog 5067-5579, 5067-5580). Illumina’s Library Prep kit (RS-122-2102) was used to generate ∼281 bp cDNA fragments, on average, which were quantified using Qubit Flex Fluorometer (catalog Q32852) and quality checked using Agilent’s TapeStation (catalog 5067-5584, 5067-5585). Next, 75 bp paired-end sequencing was performed on Illumina’s NextSeq500 platform using their high-output (150 cycles) flow cell kit (FC-404-2002) according to manufacturer’s protocol.

#### Data processing before analysis

The quality of reads was assessed using FastQC (Version 0.10.1). The adapter sequences “AGATCGGAAGAGCACACGTCTGAACTCCAGTCA”, “AGATCGGAAGAGCGTCGTGTAGGGAAAGAGTGT” were trimmed from the 3’ ends of the reads with Cutadapt from Trim Galore package (Version 0.3.7) (http://www.bioinformatics.babraham.ac.uk/projects/trim_galore/). The trimmed reads were mapped against the reference genome of *T. gondii*, GT1 (ToxoDB release 41), and assembled with HISAT2 (Version 2.0.5) [23]. The overall alignment rate was above 90% for 6-hour extracellular samples and between 25% to 60% for intracellular samples. SAM files obtained from alignment results were processed using SAMtools (Version 1.4.1) and the relative abundance of transcripts were estimated using featureCounts [24]. Counts per million (CPM) values per gene were quantified using cpm() function from the edgeR Bioconductor R package (Version 3.24.3) [24]. Genes with CPM value > 2 in at least 3 samples were retained for further analysis. Gene counts were normalized and scaled to logarithmic form using edgeR’s TMM method (trimmed mean of *M* values) with DGEList(), calcNormFactors() and cpm() functions. The cpm() parameters were: y = DGEList.obj, log=TRUE, prior.count=3, and normalized.lib.sizes=TRUE. Batch effects were examined and visualized by hierarchical clustering using the R function hclust() with the default parameters and logCPM expression values. Hierarchical clustering classified some samples in a single batch that were relatively far from their associated biological replicates based on Euclidean distance metric. This was also observed in the MDS plots (Multidimensional scaling plot of distances between gene expression profiles). The plotMDS() function from edgeR was utilized to generate the plot and visualize outliers. Batch correction was performed using removeBatchEffect() from the Limma R package with the following parameters: x = logCPMexpr, batch=batch, design=design to correct for unknown technical batch effects and avoid their ramification on downstream analysis.

#### Differential expression analysis (DEA)

DEA was carried out using edgeR. A design matrix was generated with model.matrix() function for the treatments/conditions (13 factors) and batches to perform pairwise comparisons. A normalized DEGList object was constructed from counts and treatments with DGEList() and calcNormFactors(). The estimateDisp() function was then used to estimate the dispersion based on Cox-Reid profile-adjusted likelihood (CR). The estimateDisp() parameters were: y = DGEList.obj, design = design.matrix, and robust=TRUE. The quasi-likelihood negative binomial generalized log-linear model was then fitted to the count data by glmQLFit() with robust parameter set to TRUE. The returned object of class DGEGLM from glmQLFit() was passed to glmQLTest() to ascertain the DEGs. The most DEGs ranked either by FDR adjusted p-value (q-value) or by abs(log-fold-change) were extracted with the function topTags(). Transcripts with two-fold and higher differences in their expression levels (abs(logFC) > 1) and q-value < 0.05 were considered significant DEGs.

#### Clustering analysis

Temporal patterns of the DEGs were captured using R package TCseq (Version 1.12.1)[74]; tca() function was used to generate a tca object for the time course temporal analysis. Tca() function requires 3 inputs including experiment design, genomicFeature (gtf) file, and counts table. The experiment design was generated from RNA sequencing information with columns having the sample ids and time points. The GT1 annotation file (gtf format) with the gene’s ID and location was used for genomicFeature parameter. Since tca() only accepts the raw integer counts (not normalized expression) we used the MDS plots and hierarchical clustering to identify the noisy samples (replicates) and removed them from the raw counts data. Biological replicate one from passages 35 and 210 were excluded from the temporal analysis. Once the tca.obj was constructed, Dbanalysis() quantified the log FC values using the tca.obj and by fitting negative binomial generalized linear model to the read counts. Then normalized time course table containing expression values of all extracellular samples was created with timecourseTable() function with these parameters: tca = tca.obj, value = “expression”, and filter = FALSE. Clustering of time-course data was done in an unsupervised manner using Cmeans (CM) method as implemented in TCseq in timeclust(). The total number of clusters was set to 8 (trial and error). Detected patterns were standardized and visualized. To label the detected clusters as being either strongly up- or down-trending, a linear regression model was fitted to the data in each cluster. The clusters with a large positive or negative slope of the regressed line and *R*^2^> 0.5 were selected as trending clusters, resulting in two up trending and one down trending cluster.

#### Linear mix-effect models with regression splines

Regression Analysis (RA) was performed to identify the genes that demonstrate strong correlation with evolution of GT1’s phenotypic traits over passages (P). The phenotype measurements (plaque size, invasion efficiency, and extracellular survival) were collected at 9 time points (P12, P33, P51, …, P223) with ≥3 replicates at each time point. Three biological replicates of RNA-Seq data was collected at seven passages (P7, P11, P35, P55, P85, P148 and P210) and four passages (P7, P11, P85, P148) for extracellular and intracellular parasites, respectively. A mixed-effect regression spline model was fitted to the phenotypic and RNA-Seq time course data separately using the following mixed-effect model:

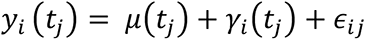

where *μ*(*t*) is the fix effect corresponding to population mean, *γ_i_*(*t*) stands for the random effect corresponding to deviation of each gene (phenotype) from mean at each time point and *ε_i,j_* is the assumed independent normally distributed noise. The random effect is assumed to be generated from a Gaussian Process *γ*_*i*_ ∼ *GP*(0, *δ*) with mean 0, which implies that the measure values of *γ_i_*(*t_j_*) are normally distributed with covariance matrix *D*(ℓ, *s*) = *δ*(*t*_ℓ_, *t_s_*). The mixed-effect model was applied through its implementation in sme() function from the R package sme [75] with time course data and criteria = ‘AIC’. The mean curve was calculated for each gene (phenotype) by sme(). Since the time points at which the RNA-seq and phenotype data were collected were not aligned, we generated a custom script to fit a natural cubic spline to the returned coefficients from sme(). Using the fitted spline, the missing values corresponding to gene expression (phenotypes) were predicted at several time points spanning the common rage of passages. Next, we calculated the “Spearman” and “Pearson” correlation between the fitted means of all genes and all phenotypes using the R function cor().

The fitted object returned by the combination of sme() and spline() models was used to visualize the mean curve along with the confidence bands at 95% level. The variability around the mean curve was derived from the variance-covariance matrix of the fitted model quantified with vcov() R function given the fitted object.

#### GLASSO TF analysis

We assembled a PPI network involving annotated *T. gondii* GT1 transcription factors (TFs) comprising all annotated ApiAP2 domain containing proteins (including an unnamed ApiAP2 on chromosome IX, TGGT1_215895, which we named for the next available numeric to AP2IX-10, and seven Myb domain containing proteins (TGGT1_200385 (BFD1), TGGT1_203950, TGGT1_213890, TGGT1_264120 (Myb1), TGGT1_275480, TGGT1_306320 (Myb2), TGGT1_321450 (Myb3)) by applying a Gaussian Graphical Model (GMM) to our RNA-seq data. The goal of GMM is to capture direct pairwise relationship between two nodes of a graph by estimating the covariance and the precision matrix Θ^-1^ from sample the covariance matrix *S*. Each AP2 and Myb represents a node in the graph and the edges represents the “direct” interaction between them after accounting for partial correlations. The objective function of the GGM is given by

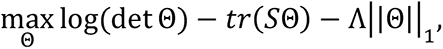

where *S*, Θ, and Λ are the empirical covariance, precision matrix and the penalty matrices, respectively.

The normalized expression values (CPM) data across all the samples and conditions was used to determine the empirical co-variance matrix. Estimation of a sparse inverse covariance matrix using a lasso *L*_1_ was fitted using GlassoFast package [76]. For selecting the right regularization parameters, Λ in estimating the inverse covariance matrix, a path of parameters (from 0.1 to 1 with 0.1 step size) was used and the Kullback–Leibler divergence was calculated. The best Λ was chosen as the value where the second derivative of the *KL*(Θ_Λ*j*_, Θ_Λ*j+1*_) function was smaller than a constant [77]. Once the optimal Λ was selected, the inverse covariance matrix was estimated accordingly. Strong associations between AP2s, were identified as those with absolute value of partial correlation greater than 0.01. The Igraph R package (V1.2.5) was used to visualize the network.

#### Ranking gene candidates

Genes were ranked based on the following criteria: 1) Significant correlation (0.7) with at least one phenotype (∼3000 genes); 2) differentially expressed in at least one passage compared with base line P11 (986 genes); 3) trending (up or down, 439 genes). This resulted in a list of 300 genes.

#### Software code

available on GitHub through: https://github.com/umbibio/ToxoplasmaGondii

### Plasmids and parasite strain generation

All oligos used are provided in Supplemental Table 7. Synthesized and annealed forward and reverse oligos, serving as sgRNAs, were cloned into *Bsa*I digested pU6-Universal plasmid (Addgene #52694) to generate our final CRISPR/Cas9 plasmids [78]. A DHFR selection cassette was amplified with 60 bp primers to yield a 2700 bp repair template containing the entire 5’UTR, 3’UTR, and CDS of DHFR, along with 39 bp arms complementary to the site of Cas9-mediated DSBs within the gene of interest (GOI). To KO a GOI, 20 μg of each CRISPR/Cas9 plasmid were co-transfected with 20 μg of the DHFR selection cassette to enabled DSB and homology-directed repair at the locus. Successful homologous recombination of the DHFR selection cassette into the GOI locus was confirmed with diagnostic PCR primer sets. Successful ablation of mRNA expression was confirmed by qRT-PCR.

### Life stage score analysis

Previously published tachyzoite, merozoite, bradyzoite and sporozoite RNA-Seq datasets were used to generate stage-scores [26, 79]; all RNA-Seq datasets were obtained from ToxoDB.org and downloaded as TGME49 gene IDs, which were then converted into TGGT1 gene IDs using the “syntenic orthologs” tool on ToxoDB.org. This process excluded 496 TGGT1 gene ID’s from the analysis. Four singular timepoints from these available datasets were chosen for DE analysis (“tachyzoites”, “tissue cysts”, “Mero 3”, and “sporozoite day 4”). For each gene, all three possible differential expression scenarios between these four datasets were calculated. For each gene, the number of *≥*2-fold upregulated scenarios were enumerated and was considered the “stage-score” (score = 0 to 3). To validate the established stage scores, three previously published gene sets were examined for significant enrichment in their indicated life stage [28]. For analysis of our RNA-Seq data sets, the DE-score of our upregulated and downregulated gene sets were individually calculated.

#### Bootstrap analysis

To empirically estimate the statistical significance of the scores we performed bootstrap analysis as follows: For each gene set 1000 random genes were sampled with replacement and the mean score of all four life stages were calculated. The distribution of the mean scores for each life stage were estimated from the bootstrap samples and the *p*-value of the observed mean score of each gene set was calculated by summing over the right-tail of the distribution.

### Enrichment analyses

Previously published gene sets [28] along with tachyzoite were utilized for GSEA. Fisher’s Exact Test was used to assess the statistical significance of the overlaps between differentially expressed genes in various contrasts, trending genes and classes derived from our data and the gene sets. The R function fisher.test() calculated p-values corresponding to the overlaps between two gene sets. The fisher.test() parameters were as following: x = contingency.table, alternative = “greater”. P-value ≤ 0.05 represent significant enrichment of the gene set in the priori database. GOEA and MPEA was performed using the “Analyze results” feature on ToxoDB.org (https://toxodb.org/toxo/analysisTools.jsp) [80].

### Protein modelling

Protein Homology/analogy Recognition Engine v2.0 (Phyre2) prediction and analysis tools (http://www.sbg.bio.ic.ac.uk/~phyre2/html/page.cgi?id=index). Subsequently, we predicted the impact of the SNPs on the structure using Missense3D predictive structural analysis (http://www.sbg.bio.ic.ac.uk/~missense3d/) [21].

### Statistical analysis

A student’s two-tailed equal variance *t* test was used to determine the significance (*p*-value) of evolved samples compared to the lowest passage (P) sample. Adjusted *p*-values (*q-*values) were calculated using the false discovery rate (FDR) method. For stage-score analysis, both a student’s two-tailed *t* test and an independent bootstrap analysis (n=1000 random sampling) was used to determine significance. A one-sided Fisher’s exact test was used to quantify the significance (*p-value*) of each gene set’s enrichment in the priori database.

## Acknowledgements

We thank Dr. Karen Zhu and Sandra Dedrick at BC’s Sequencing Facility, Drs. Klemens Engelberg, Daniel Tagoe, Michelle Meyer, Babak Momeni, and Tim van Opijnen for their intellectual support, Emily Stoneburner, William Pisano, Humza Rashid and Jingjing Lou for technical support and Dr. Mahipal Singh for sharing reagents.

## Funding information

This work was supported by grants AI081220, AI150090 and AI122923 from the National Institutes of Health.

## Competing interests

The authors declare no competing interests.

## Author contributions

All parasite phenotype data was generated, analyzed, and interpreted by VAP and MJG. All NGS data was generated by VAP and analyzed by AF, YR and AV under the supervision of GTM and KZ, respectively. Various enrichment and life-stage analyses were generated by VAP and YR, under the supervision of MJG and KZ, respectively. Statistical analysis was performed by YR and KZ. Re construction of AP2 network with GGM was performed by AV.

## Data availability

Short read DNAseq and RNAseq data are deposited as FASTQ data to the NCBI Sequence Read Archive (SRA) with submission number 9045827.

## Supplementary Material

### Headers to Supplementary Tables

**Supplementary Table 1. 12 DEGs identified in intracellular GT1 parasites during lab-adaptation.**

RNA-Seq of intracellular B2 GT1 populations at passages P11, P84, P148, followed by DEA, identified 12 significant DEGs (FC *≥*2, q-val *≤* 0.05), relative to the lowest passage, P11. Log_2_ fold change is show; red text indicates statistical significance (q-val *≤* 0.05).

**Supplementary Table 2. 986 DEGs identified in 6-hour extracellular GT1 parasites during lab-adaptation.**

RNA-Seq of 6-hr extracellular B2 GT1 populations at passages P11, P35, P55, P85, P148, and P210, followed by DEA, identified 986 significant DEGs (FC *≥*2, q-val *≤* 0.05), relative to the lowest passage, P11. Of those, 435 DEGs were upregulated and 551 DEGs were downregulated. Log_2_ fold change is show; red text indicates statistical significance (q-val *≤* 0.05).

**Supplementary Table 3. GSEA, GOEA, and MPEA of the 986 DEGs and 300 DEGs identified in 6-hour extracellular GT1 parasites during lab-adaptation.**

GSEA, GOEA, and MPEA was performed on upregulated and downregulated DEG identified in Supplementary Table 2 (986 DEGs) and Supplementary Table 4 (300 DEGs). GOEA and MPEA results were obtained from ToxoDB.org.

**Supplementary Table 4. 300 DEGs identified in 6-hour extracellular GT1 parasites during lab-adaptation.**

Refined list (derived from Supplementary Table 2) of the highest phenotype correlating (*R*^2^*≥*0.70, Spearman correlation) DEGs, as identified by the overlap of DEGs identified by DEA, TC-seq, and RS analysis. Of the 300 DEGs, TC-seq identified 3 main expression trends (i.e. clusters) identified as clusters 1, 2, and 6. Cluster 1 consists of 119 upregulated DEGs; cluster 2 consists of 107 downregulated DEGs; and cluster 6 consists of 74 upregulated DEGs. The Spearman correlation coefficient of all 300 DEGs is provided.

**Supplementary Table 5. 300 DEGs subcellular localizations.**

HyperLOPIT subcellular localization data available for tachyzoites [30] was downloaded for ToxoDB and in case of discrepancy between the statistical methods, manually curated where possible, including assignment of two transcription factors to the nuclear pool. The ‘mixed’ pool refers to where a call could not be made on the conflicting statistical assignments.

**Supplementary Table 6. 2393 DEGs identified between intracellular and 6-hour extracellular GT1 parasites.**

RNA-Seq of intracellular and 6-hr extracellular RH and GT1 (B0-P7) parasites, followed by DEA, identified 2393 DEGs that are significantly (FC *≥*2, q-val *≤* 0.05) different between the intracellular and extracellular state in both RH and GT1 (B0-P7) parasites. Of those, 1189 DEGs are upregulated and 1204 DEGs are downregulated. Other GT1 passages are also shown for comparison. Log_2_ fold change is show; red text indicates statistical significance (q-val *≤* 0.05).

**Supplementary Table 7. Oligonucleotide sequences used.** Table provided indicates oligo sequence used for: M13 primers used for PCR-Sanger sequencing of known RH and GT1 SNPs and for genotyping the P4-flippase (TGGT1_2455510) SNPs; protospacer sequences used to generate KOs; primers used to generate DHFR cassette used as a repair template; diagnostic PCR primers used to confirm genomic integration of DHFR cassette; and qRT-PCR primers used to confirm mRNA ablation in KOs.

**Source Data.** Contains source data for Figure 1c, Figure 1d, Figure 2a, Figure 2b, Figure 2c, Figure 2d, Figure 5a, Figure 7e, Figure 7f, Supplementary Figure 1a, Supplementary Figure 1b, Supplementary Figure 1c, Supplementary Figure 1d, Supplementary Figure 4, Supplementary Figure 6d, Supplementary Figure 7, Supplementary Figure 8, Supplementary Figure 9a, and Supplementary Figure 9b.

**Supplemental Figure 1.**
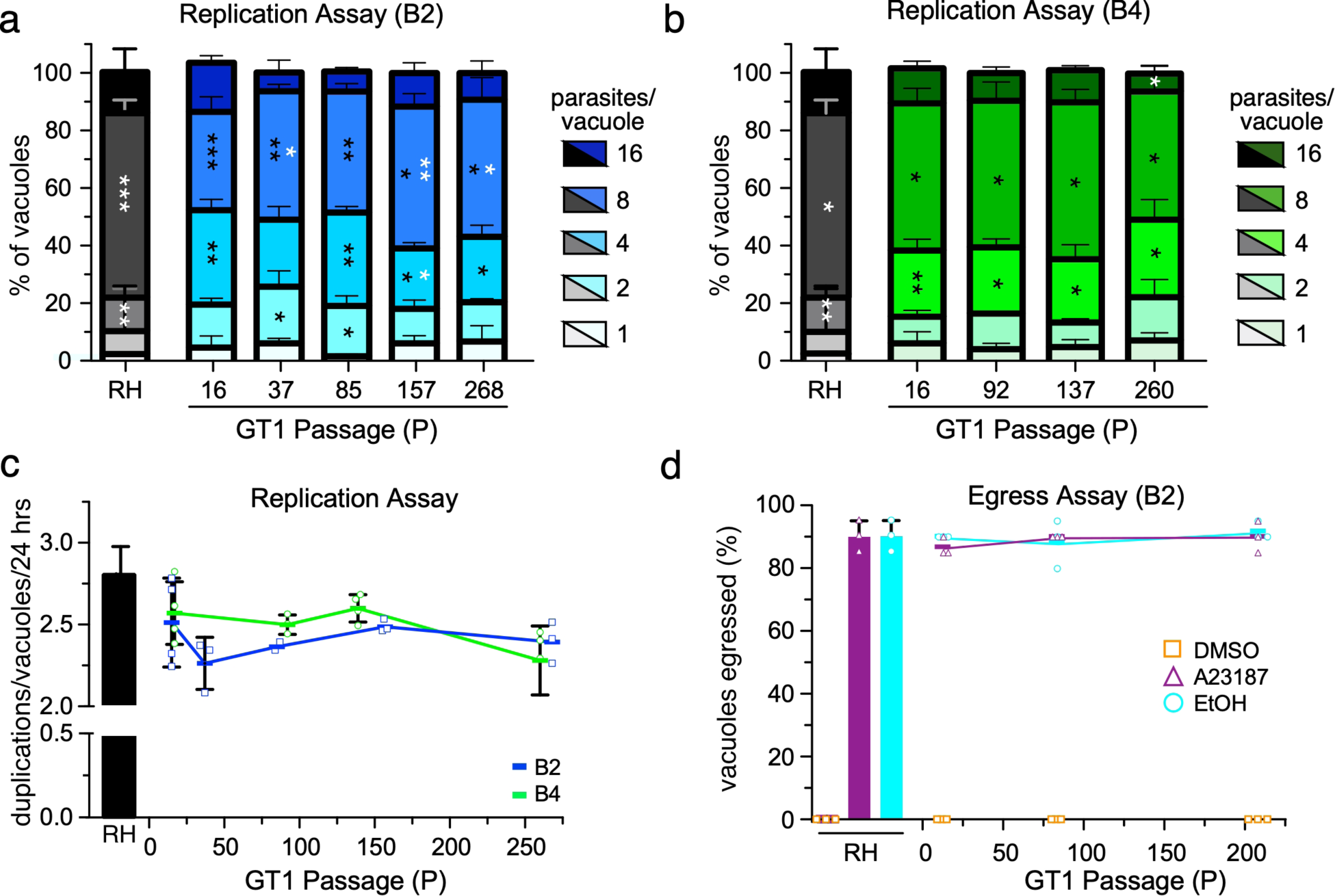
Virulence traits of the intracellular milieu are not affected by lab-adaptation. **a** Replication assay of RH and GT1 populations B2 or **b** B4 parasites at the indicated timepoints. The number of parasites per vacuole in 100 random vacuoles were quantified 24-hours post infection by IFA. Mean of *≥*3 biological replicates with SD is plotted. Black asterisks (*) indicate the *p-*value of the indicated GT1 passages relative to RH; white asterisks indicate *p-* value of the indicated GT1 passage relative to the respective population’s earliest passage (GT1-P16); * = *p*-value *≤* 0.05, ** = *p*-value *≤* 0.01, *** = *p-*value *≤* 0.001. **c** Same dataset as in a, b represented as average number of parasite duplications during the first 24 hours of inoculation. Colored blocks indicate mean of *≥*3 biological replicates for B2 and B4 (shown as circles and squares, respectively) with sizeable error bars representing SD. **d** 30-hour intracellular parasites were treated for five minutes with either control (DMSO), calcium ionophore (A23187), or ethanol (EtOH). The number of egressed and non-egressed vacuoles were enumerated by IFA. Mean of 3 biological replicates (shown as squares, circles, and triangles) with sizeable SD is plotted.

**Supplemental Figure 2.**
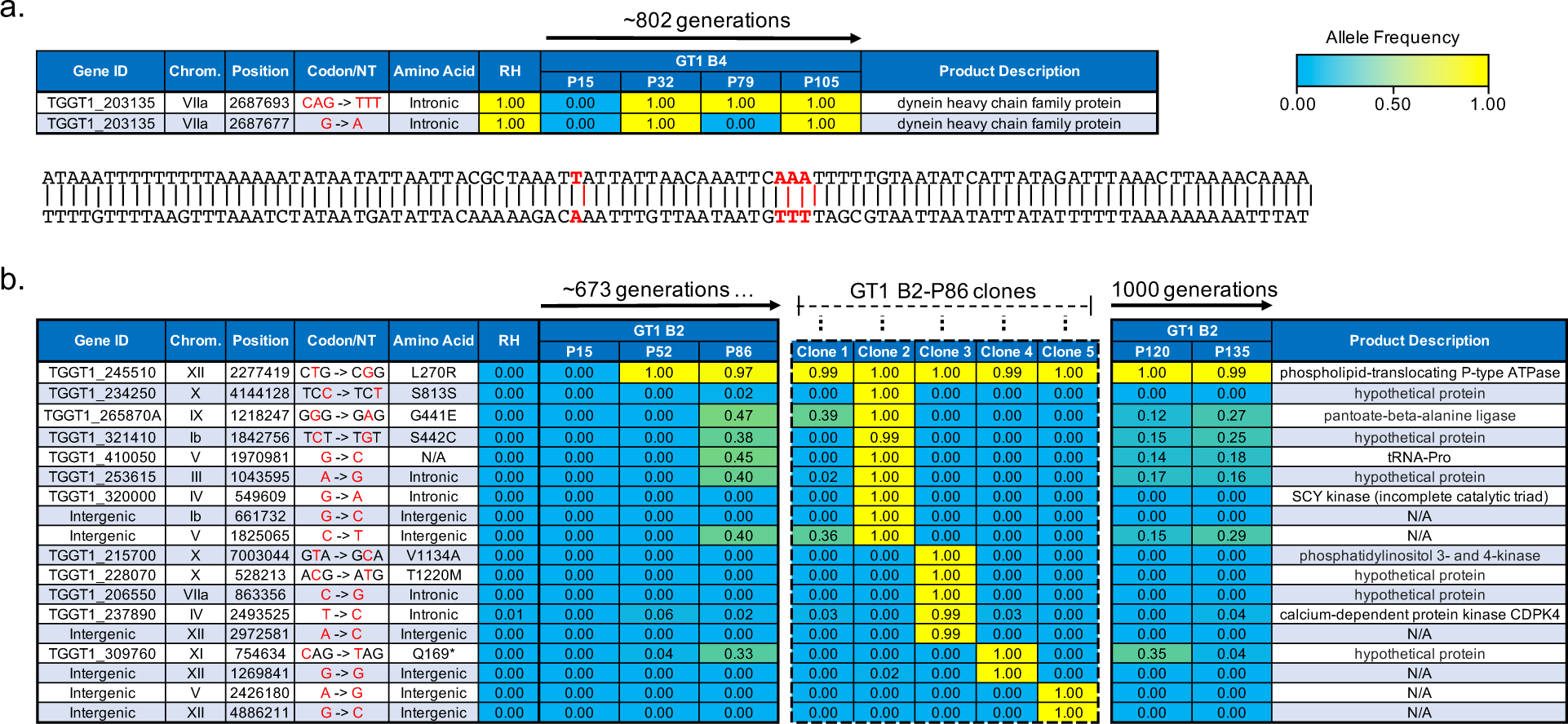
Mutations arise, but rarely fix, during the first 1000 generations of i*n vitro* lab-adaptation. WGS identified the emergence of indicated mutations with the indicated allele frequency in GT1 populations; RH is shown for reference. **a** Mutations identified in GT1 population B4 during >800 generations of lab-adaptation; 100 bp nucleotide sequence surrounding the site of mutations is shown below. **b.)** Mutations identified in GT1 population B2, along with five B2-P86-derived clones. N/A indicates the gene does not result in a protein product. For all panels: allele frequency represents the percentage of reads with the indicated allele.

**Supplemental Figure 3.**
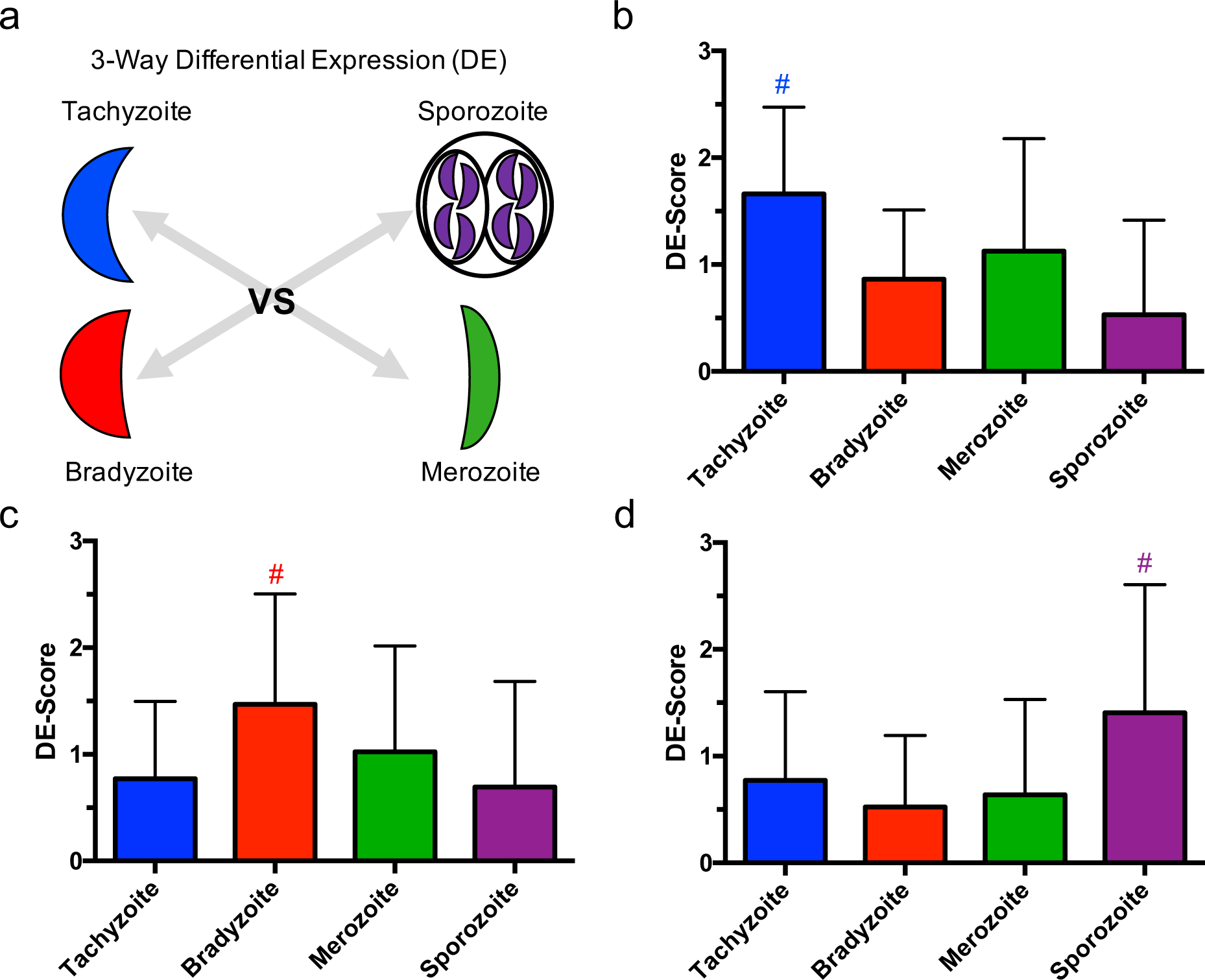
Developing and validating *T. gondii’s* life stage score analysis. **a** Available tachyzoite, bradyzoite, merozoite, and sporozoite RNA-Seq data sets (ToxoDB.org, see methods) [25–27] were utilized for differential expression analysis (DEA). Each stage was compared to the three other stages (e.g. tachyzoite vs. bradyzoite; tachyzoite vs. sporozoite; tachyzoite vs. merozoite). The number of times each gene was significantly upregulated in this 3-way DEA was enumerated to yield a maximum score of 3 or minimum score of 0 for each gene. The average enumeration of previously published lists [28] of genes was calculated for **b** tachyzoite-, **c** bradyzoite-, and **d** sporozoite-associated genes. Color-coded hashtags (#) indicate *p-*value *≤* 0.05, as determined by an independent bootstrap analysis (n=1000 random sampling) for each individual life stage. Error bars indicated SD.

**Supplementary Figure 4.**
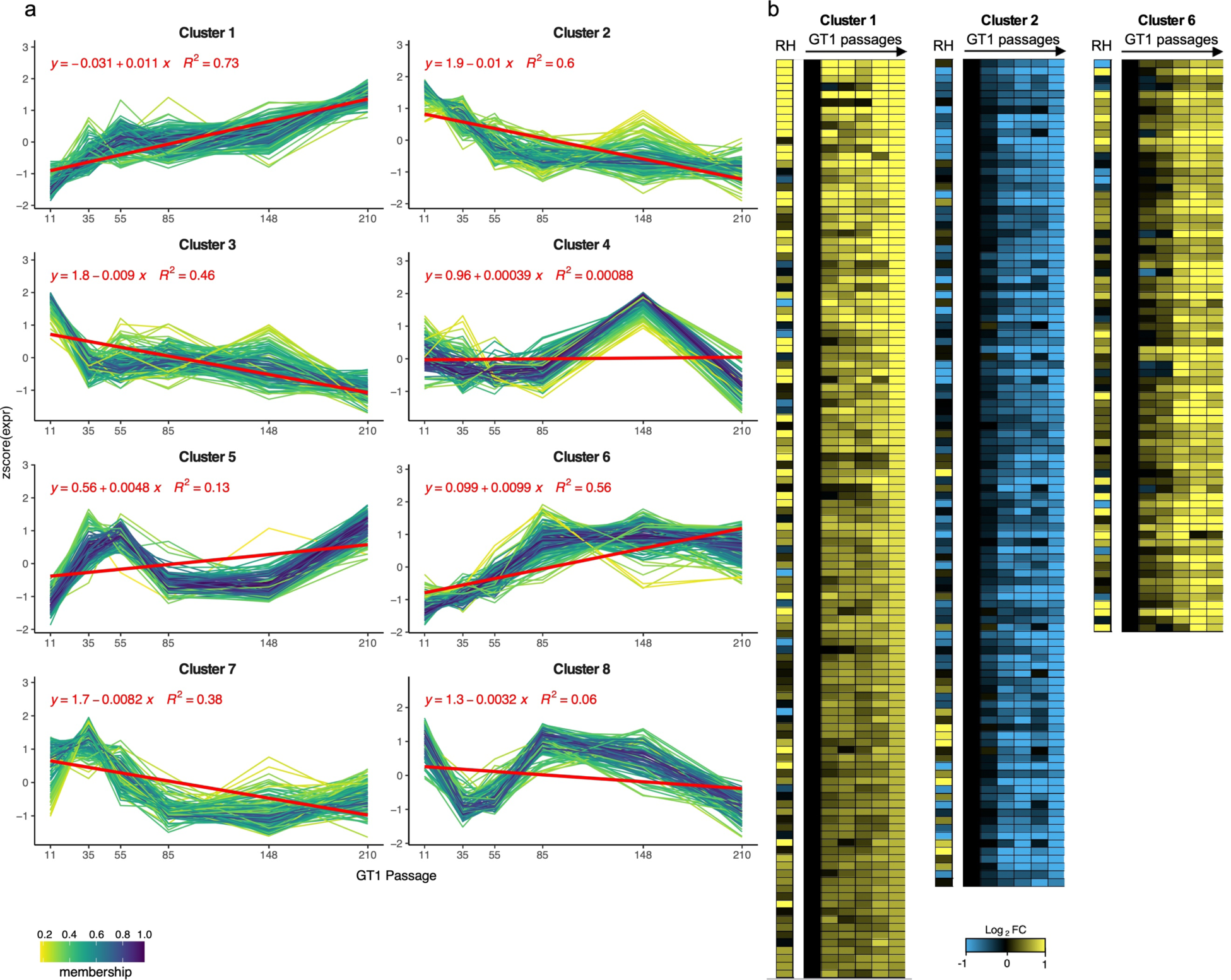
Up and down trending gene clusters identified by TCseq. **a** Normalized expression of all identified gene clusters. The genes comprising the 300 candidate DEGs identified by DEA, TCseq, and RA are contained in clusters 1, 2, and 6, since only these 3 show the most linear up- or down-trends across all time points, consistent with the steady developments in phenotypes. **b** Heatmaps of the 439 genes total genes in clusters 1, 2, and 6. This gene set was intersected with the phenotype-conferring score and resulted in the selected set of 300 genes.

**Supplementary Figure 5.**
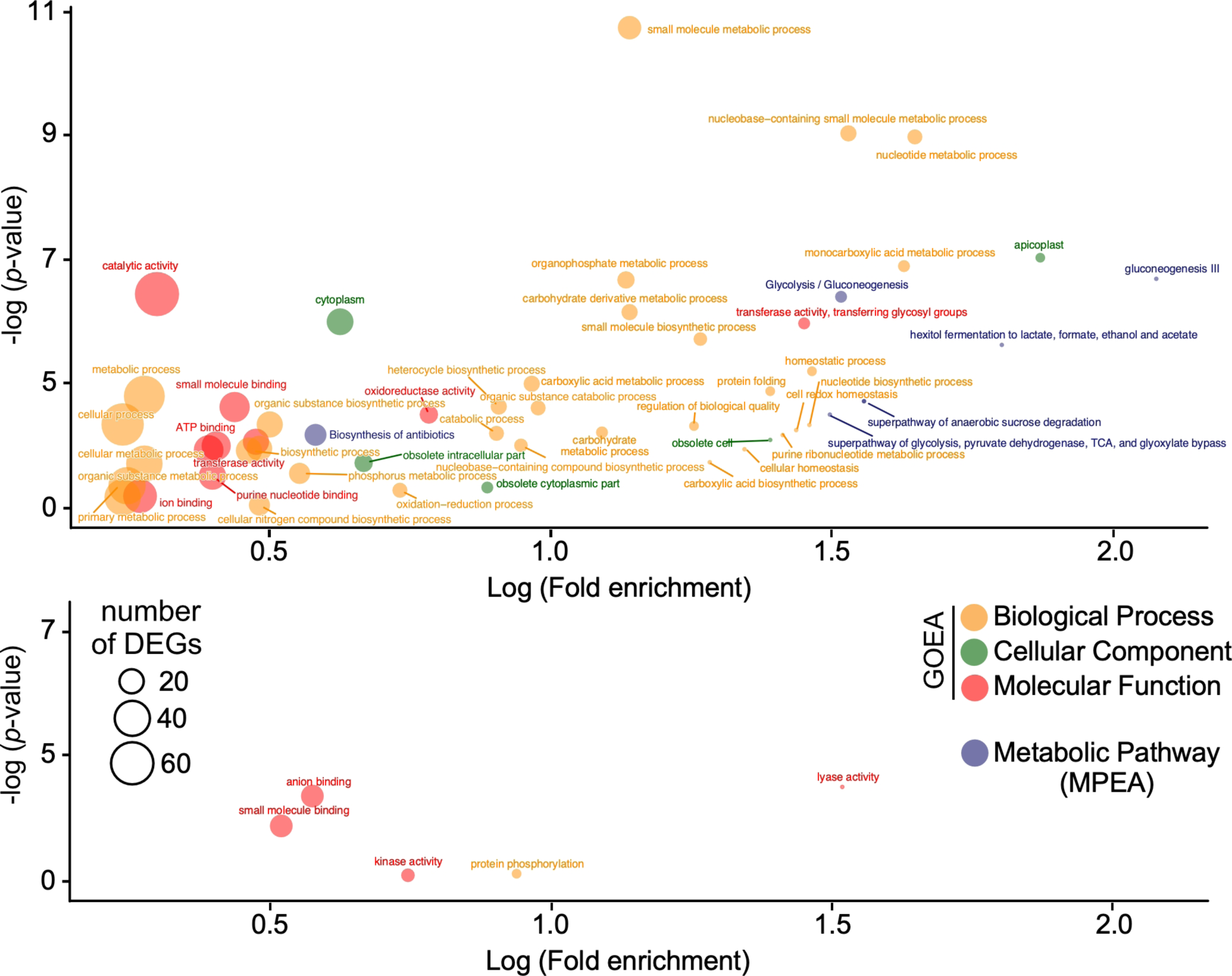
Gene Ontology Enrichment Analysis (GOEA) for the 300 most trending genes. Top panel: up trending; bottom panel: down trending.

**Supplemental Figure 6.**
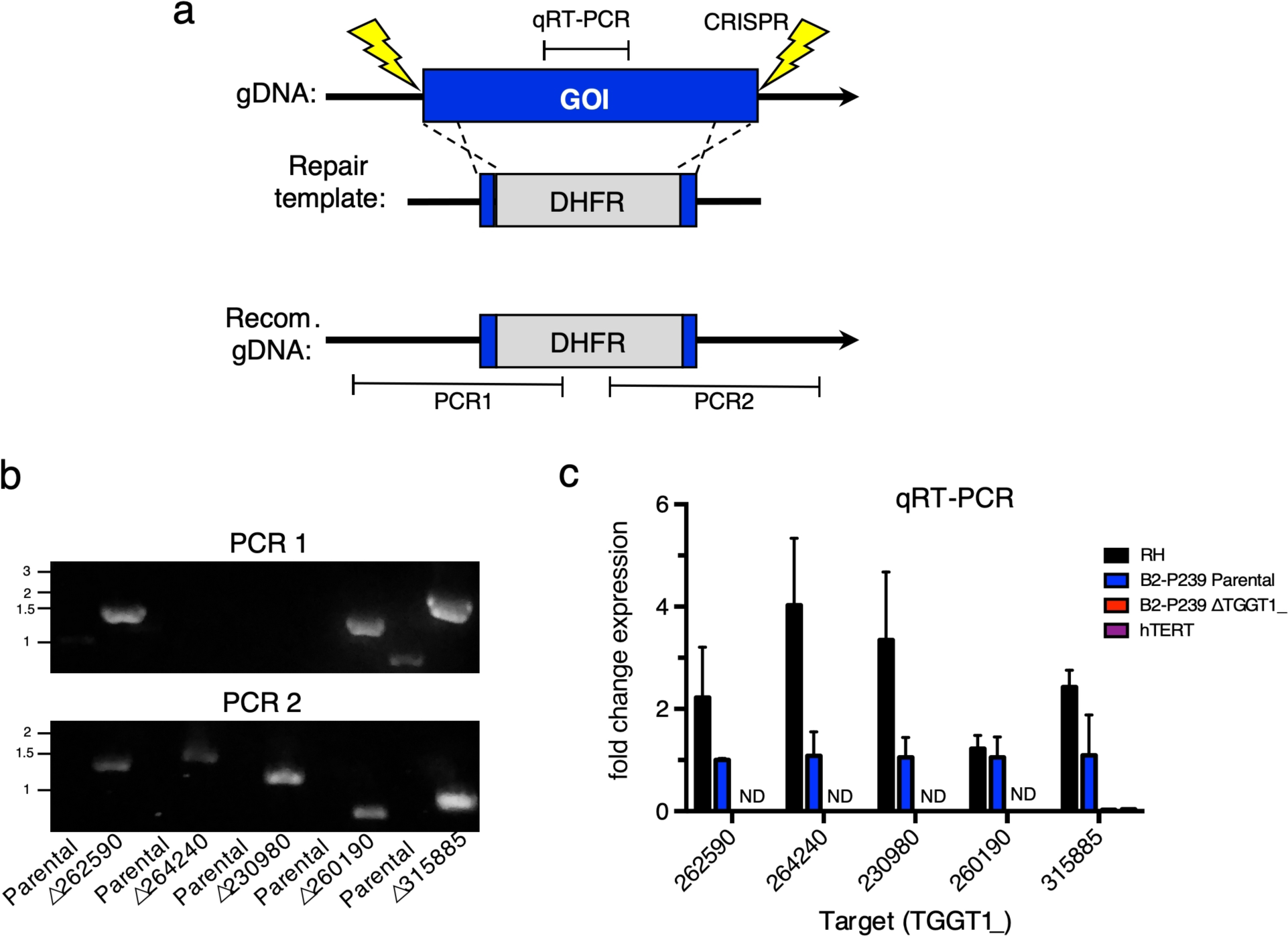
Generating genetic knockouts of regression analysis candidates by CRISPR/Cas9. **a** Strategy for generating KO of candidate DEGs. Transfection of one or two CRISPR plasmids generates a double strand break, allowing for recombination of a co-transfected DHFR selection cassette with short homologous flanks, into the genetic locus and disrupting the gene of interest. Note, sites of recombination are not drawn to scale; relative regions for diagnostic PCR to confirm integration and qRT-PCR to confirm ablation of mRNA expression are shown. **b** Diagnostic PCR of clonal KO parasites confirms integration of the DHFR cassette into the gene of interest. **c** qRT-PCR of clonal KO parasites confirms ablation of mRNA expression of the gene of interest. Fold change, relative to parental line (B2-P239) with SD is shown; N.D. indicates the expression was not detected.

**Supplemental Figure 7.**
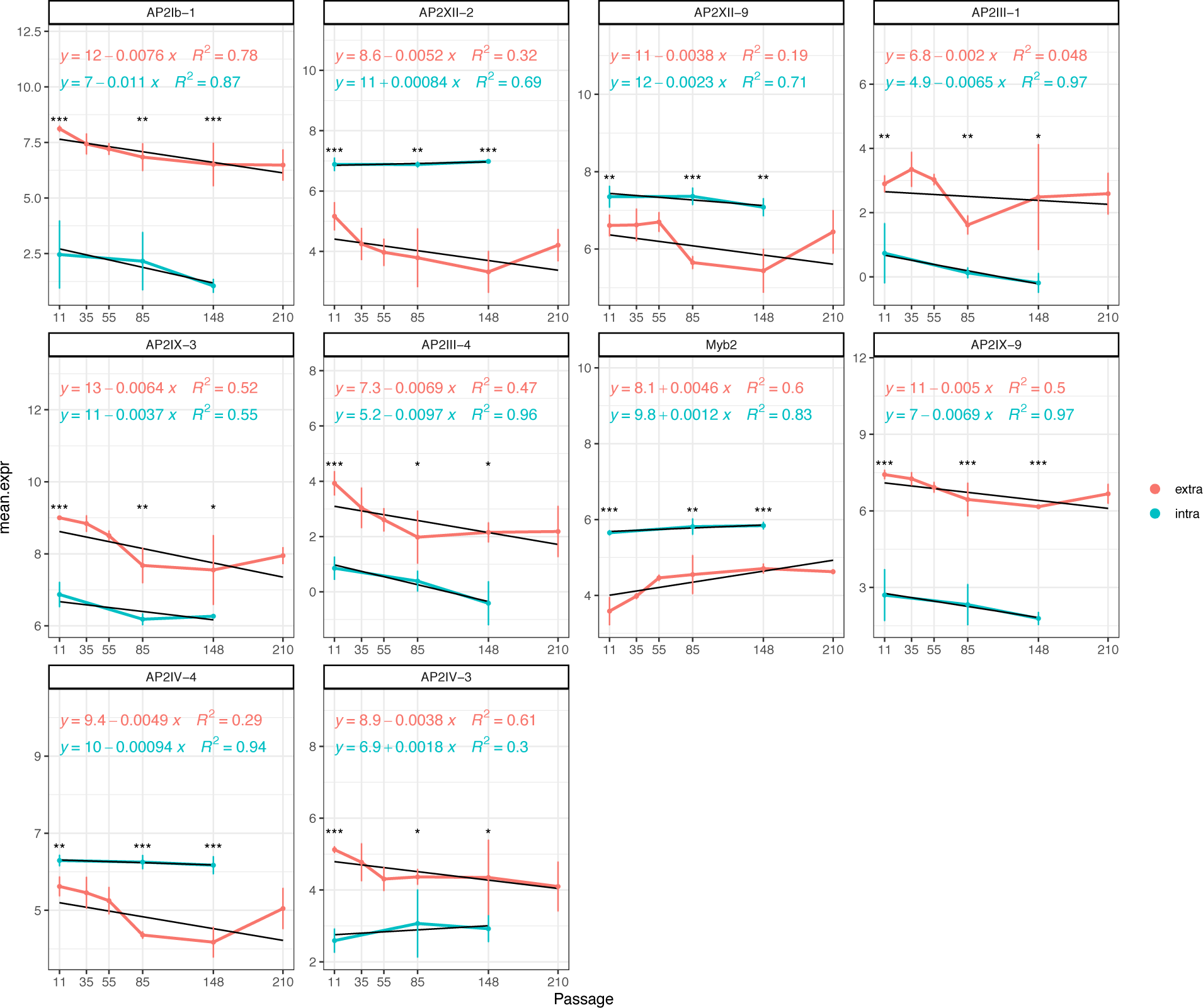
Expression pattern of 9 AP2 and 1 Myb TFs for both intracellular and extracellular parasites over the time. These TFs are significantly differentially expressed (FC *≥*2, q-val *≤* 0.05) in at least one of the passages relative to other passages in extracellular parasites. Asterisks (*) indicate the q-value of differentiated expression level between extracellular and intracellular parasites at the same passage. * = q-value < 0.05, ** = q-value < 0.005, *** = q-value < 0.0005. In each figure the black line corresponds to the linear regression line fitted to each TF’s expression value to indicate the significance of trend over time. The goodness (R^2^) and linear equation of the fit are shown.

**Supplementary Figure 8.**
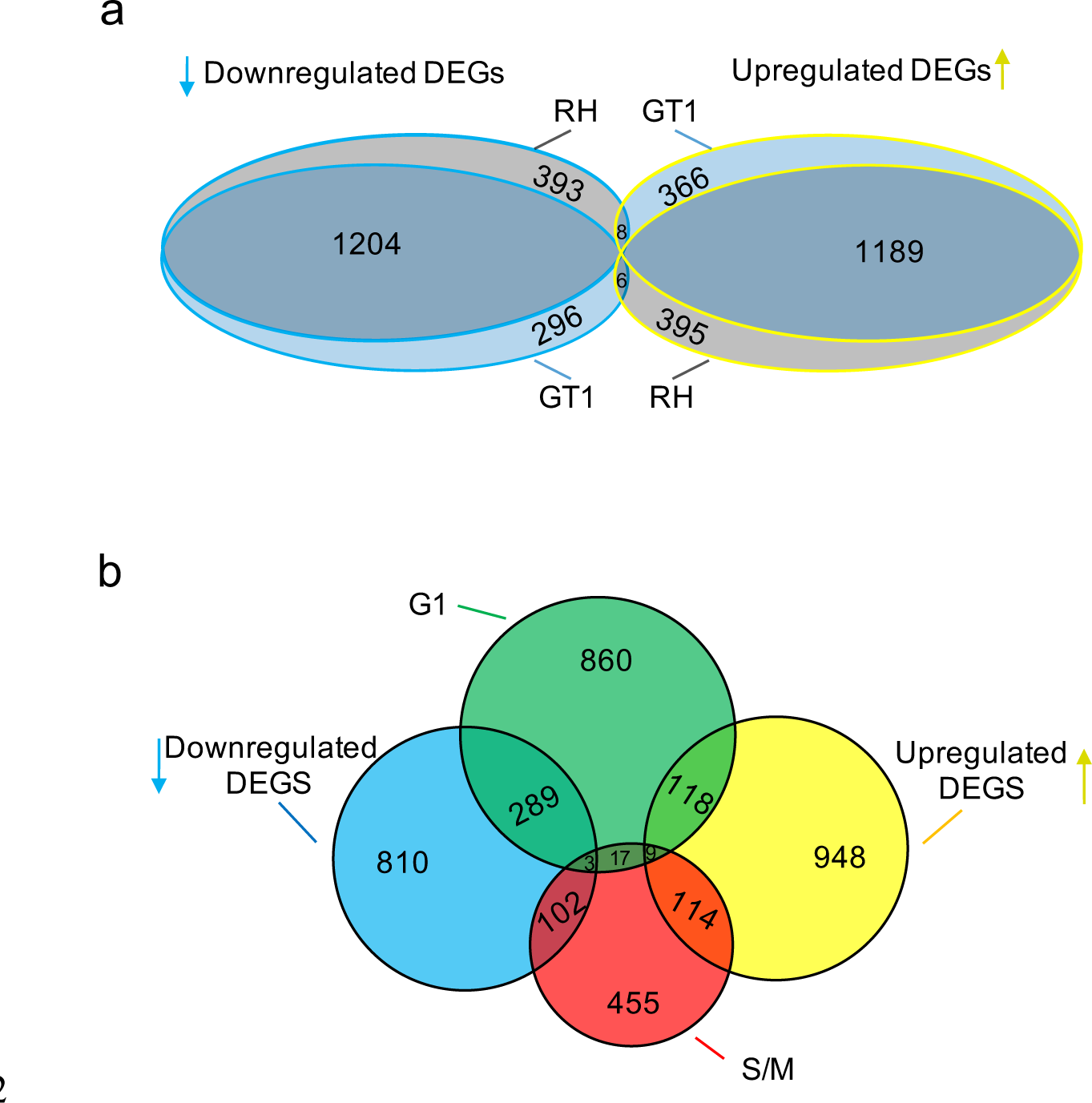
The extracellular milieu is a unique transcriptional state with many DEGs not related to the intracellular replication cycle. **a** Venn diagram of DEGs in our data sets identified between intracellular and extracellular RH and GT1 (B0-P7) tachyzoites. **b** Venn diagram of DEGs identified between intracellular and extracellular tachyzoites, as well as the two cycling sub-transcriptomes in intracellular replicating parasites [63].

**Supplementary Figure 9.**
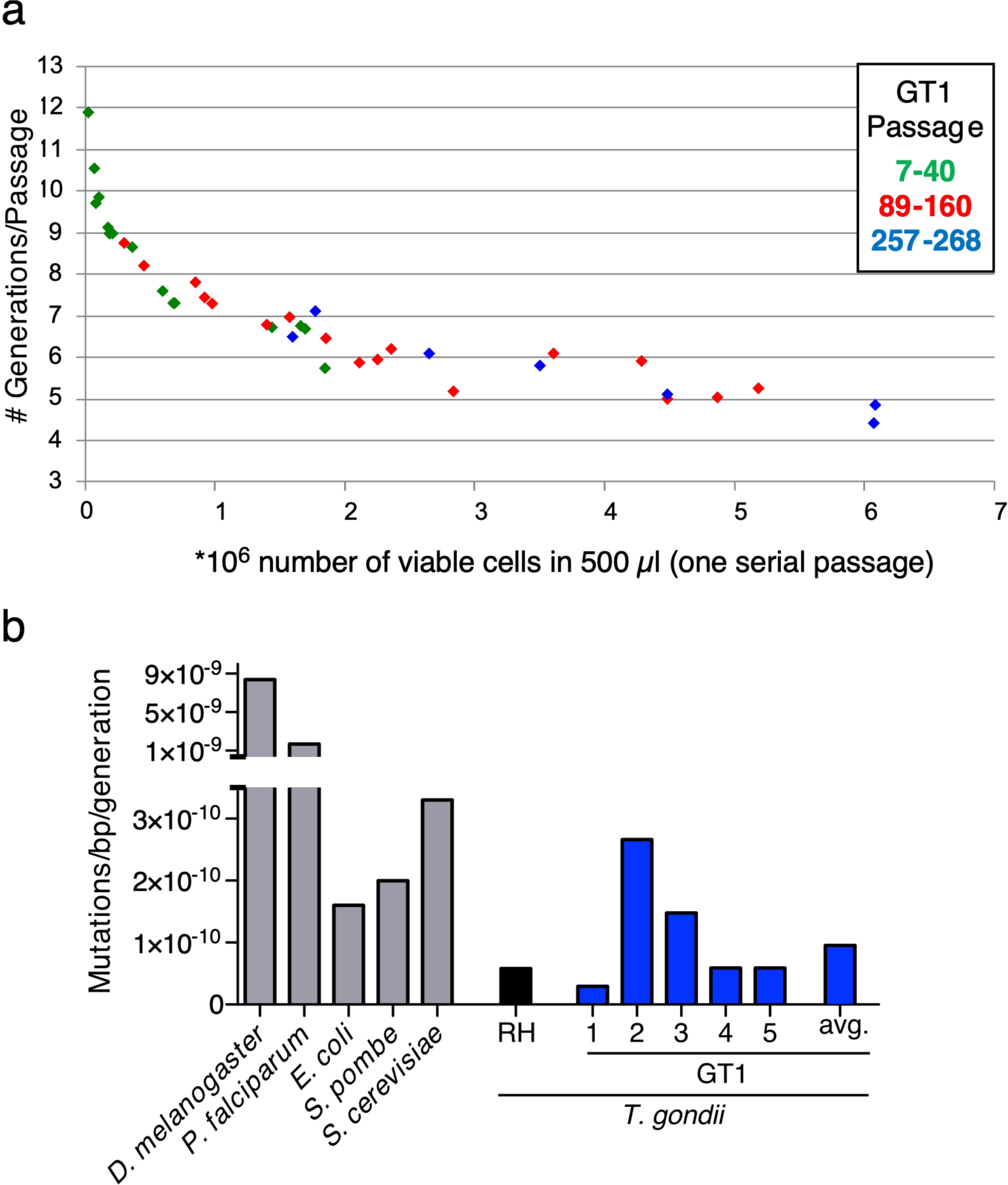
Estimating the mutation rate in the *T. gondii* GT1 strain. **a** Plaque assays and cell counts before and after serial passaging were used to determine the number of viable cells and generations/passage, respectively. The number of generations *T. gondii* will undergo per passage correlates with the number of viable cells that are passed, which is also dependent on how lab-adapted the population is. **b** Published mutation rates for *Drosophila melanogaster* [85], *Plasmodium falciparum* [86], *Escherichia coli* [14], *Schizosaccharomyces pombe* [87], *Saccharomyces cerevisiae* [88], *T. gondii* RH strain [68], and our calculated mutation rates for GT1.

## Notes

### Competing Interest Statement

The authors have declared no competing interest.

